# Evidence of maternal resilience in two mouse strains in the context of permanent mouse breeding strategies

**DOI:** 10.64898/2026.06.15.731812

**Authors:** Andrea S. Leuthardt, Camille Calmbach, Kaya Walo, Nadja Prebianca, Gaia Serra, Sander M. Botter, Rupert Palme, Paulin Jirkof, Bernadetta Tarigan, Christina N. Boyle

**Affiliations:** Institute of Veterinary Physiology, Vetsuisse Faculty, University of Zurich, Zurich, Switzerland; Swiss Center for Musculoskeletal Biobanking SCMB, Balgrist Campus AG, Zurich, Switzerland; Unit of Pharmacology and Toxicology, Department of Biomedical Sciences and Pathobiology, University of Veterinary Medicine, Vienna, Austria; Office for Animal Welfare and 3Rs, University of Zurich, Zurich, Switzerland; Applied Statistics, Department of Mathematical Modeling and Machine Learning (DM3L), University of Zurich, Zurich, Switzerland

## Abstract

Breeding female mice represent an essential but often overlooked workforce sustaining biomedical research. Despite their central role, the physiological and behavioral consequences of repeated reproductive cycles have been poorly characterized, in part because breeding animals fall outside the primary focus of laboratory animal welfare efforts, and in part because meaningful welfare readouts for laboratory rodents remain an active area of research. Here we report findings from an exploratory phenotypic study designed to capture a composite picture of maternal health in female mice of two commonly used inbred strains, BALB/cByJ and C57BL/6J, exposed to one, two, or four consecutive cycles of pregnancy and lactation, with age-matched virgin females as controls. Assessments were conducted during the final lactation period and in the five weeks following weaning, spanning behavioral, metabolic, and physiological readouts selected for their known sensitivity to reproductive or environmental challenge. Repeated reproduction altered maternal physiology, most clearly in bone microstructure, which showed progressive and dose-dependent changes across parity levels, and more subtly in body mass, energy balance, and glucose homeostasis. Behavioral readouts of maternal motivation, by contrast, remained largely stable across reproductive load. Strain differences were pervasive, underscoring that reproductive adaptation is not uniform across standard laboratory models and cautioning against generalizing from a single strain. Together, the data suggest that mouse dams demonstrate considerable resilience under intensive breeding conditions, while also highlighting that breeding shapes the maternal body in ways that accumulate across reproductive cycles and deserve greater scientific attention.

## Introduction

Rodents have been instrumental to our understanding of basic biological concepts, contributing to biomedical discoveries and drug development across disciplines to the benefit of both human and animal patients. Mice are the most widely used laboratory animal species worldwide. In 2022, around 4 million mice were used in the European Union, accounting for approximately 50% of all animals used in experimental procedures [1]. Each of these experimental mice were born to a breeding female mouse, or dam, representing an unspoken workforce sustaining biomedical research. Yet despite their tremendous contribution, the health and welfare of these breeding females have received remarkably little scientific examination.

Pregnancy and lactation are energetically demanding periods in the life of female mammals [2,3]. A key feature of rodent reproductive biology is the postpartum estrus, which occurs 14 to 24 hours after parturition, followed by lactation-induced infertility until weaning [4,5]. In permanently paired breeders, this allows females to undergo multiple rounds of concurrent pregnancy and lactation, producing a new litter approximately every three weeks. While exploiting postpartum estrus can maximize productivity and efficiency in energy-rich environments, it is disputed whether it creates energetic conflicts between suckling offspring and a simultaneous pregnancy [6,7], and the effects on maternal health are not fully understood.

While high rates of reproductive output may be interpreted as a proxy for overall fitness, and the apparent capacity of dams to sustain this tempo might suggest they are physiologically well-equipped to do so, observations in a more naturalistic settings tell a different story. Free-living house mice do not consistently exploit postpartum estrus, even in a resource-rich environment where food is available ad libitum. Tracking the reproductive strategies of mice in a semi-naturalistic setting, König and colleagues observed that female mice produce an average of three litters over their lifetime, with the first litter born at around 6.5 months of age and a subsequent litter following 67 days later [8]. This stands in contrast to permanently paired laboratory breeding schemes, in which a female can produce a new litter every three to four weeks. Further, recent data suggest that while reproductive experience does not shorten the lifespan in mouse dams, it can increase the immediate risk of mortality [9], which is consistent with the notion that reproductive load is not without cost.

Whether this intensive breeding schedule impacts maternal health has not been systematically investigated. In part, this reflects a structural gap in how laboratory animal welfare is approached. Because breeding animals are not experimental subjects, they often fall just beyond the scope of formal welfare concern. This is illustrated by a study that investigated the impact of breeding strategies, but primarily focused on offspring outcomes [10]. Efforts to address this gap are also complicated by the fact that defining meaningful welfare readouts for laboratory rodents remains an active and evolving area of research [10,11]. Importantly, while maternal physiological adaptations are expected, and indeed necessary, to maintain pregnancy and lactation [3], it remains unknown if these adaptations become insufficient or begin to impinge on maternal welfare under heavy and repeated reproductive demands of permanent breeding strategies.

With this in mind, and in the absence of established welfare benchmarks for breeding animals, we selected a panel of readouts for which there is existing evidence of sensitivity to reproductive, social, or environmental challenge. Maternal behavior in rodents is generally robust, yet specific components, such as pup retrieval, are sensitive to social or environmental stressors [12,13]. Similarly, exploratory behavior in elevated plus maze, light-dark box, or open field assays is suppressed under conditions of acute or chronic stress [13–15], providing a behavioral index of affective state. Fecal corticosterone metabolites (FCMs) offer a non-invasive measure of HPA axis activity that is widely used as an index of physiological stress in laboratory rodents [16–18]. At the physiological level, reproductive experience has been shown to reduce bone mineral density [19] and to deplete hepatic iron reserves [20], while also modifying metabolic health by increasing body mass and susceptibility to obesity following weaning [21].

To address this gap, we performed an exploratory phenotypic assessment of female mice exposed to increasing reproductive load, comparing dams that had undergone one, two, or four consecutive cycles of pregnancy and lactation with age-matched nulliparous control mice. Two commonly used inbred mouse strains, BALB/cByJ and C57BL/6J, were included to capture a broader range of biological variability. Assessments were conducted during the final lactation period and following weaning, with the aim of defining how repeated reproduction shapes maternal physiology and wellbeing while demands are still acute and while early recovery ensues.

## Results

### Reproductive performance outcomes

While mice were randomly assigned to one of four reproductive conditions, the rate of achieving the assigned number of pregnancies varied by condition and strain, indicating a level of self-selection toward high breeding performance, most notable in the group with four cycles. A comprehensive overview of maternal reproductive performance and offspring outcomes is provided in Table 1 and Table S1. Across both strains, the proportion of females completing successive cycles declined markedly with increasing parity. In BALB/cByJ dams, 66.7% (N = 10/15) completed one cycle, 50% (N = 9/18) completed two consecutive cycles, and 14.3% (N = 5/35) completed four cycles. In C57BL/6J mice, 100% (N = 10/10), 62.5% (N = 10/16), and 22.6% (N = 7/31) of dams completed one, two, and four cycles. Thus, the proportion of dams completing four cycles was substantially reduced in both strains.

**Table 1.**
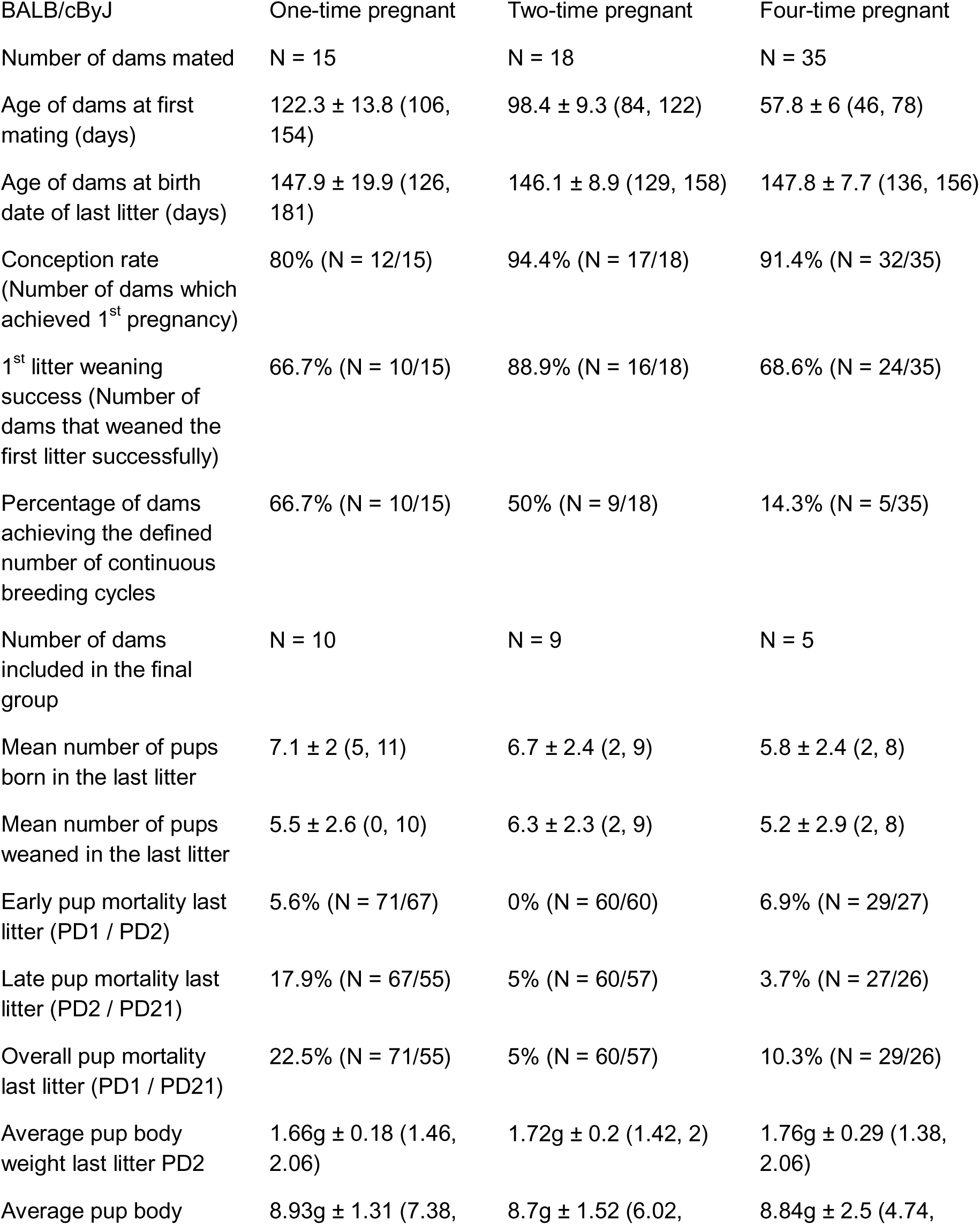
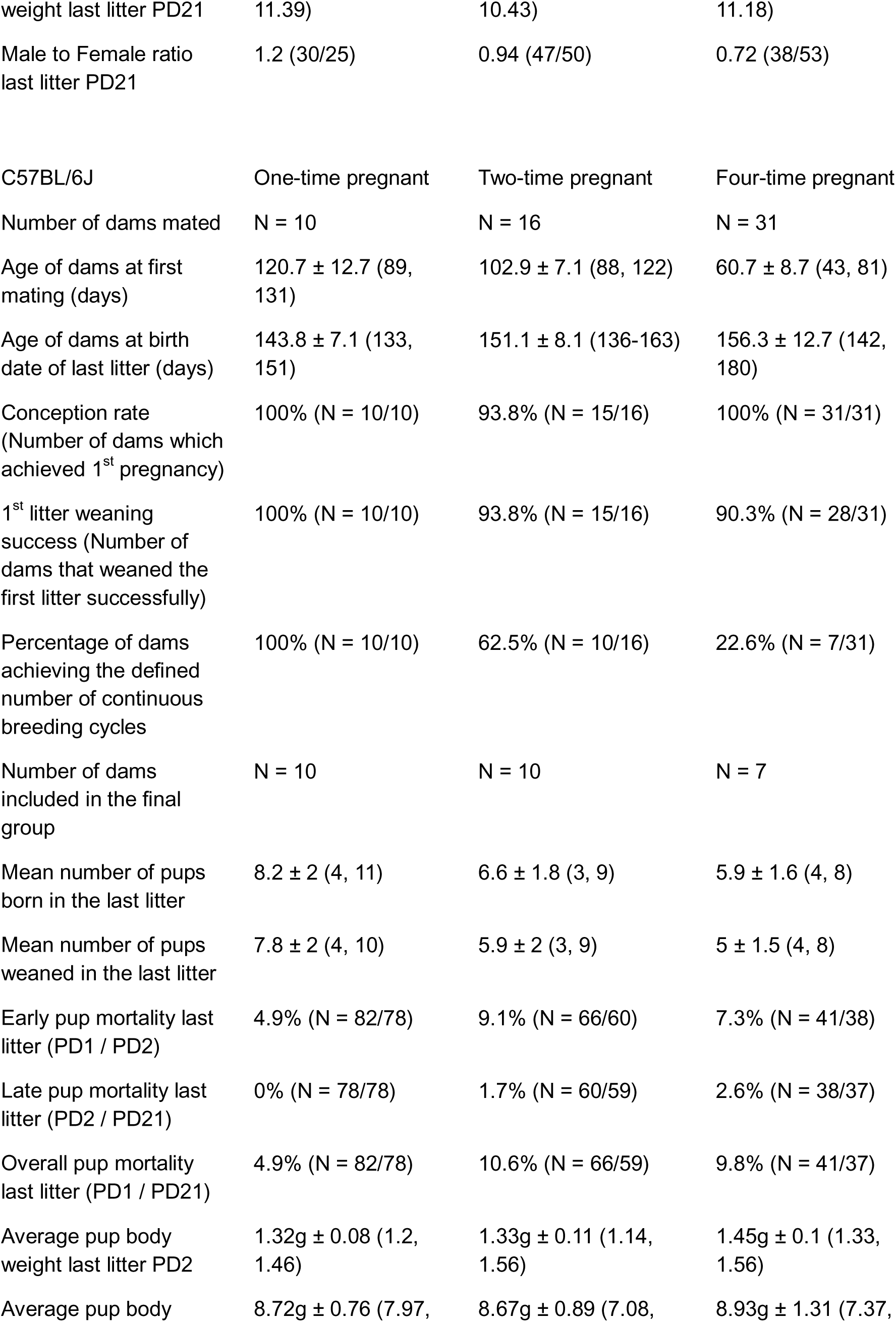
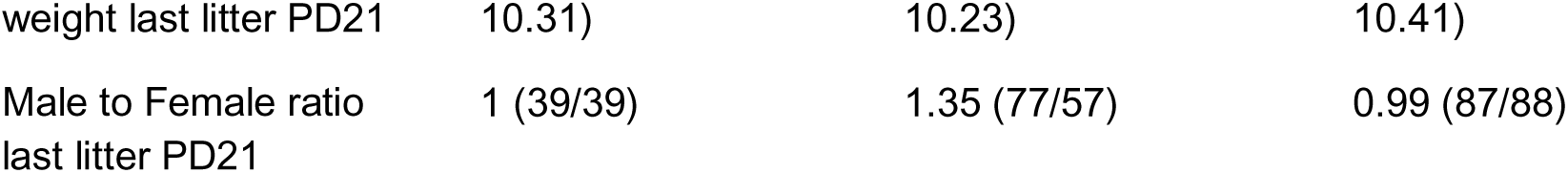
Reproductive performance and offspring outcomes of BALB/cBJ and C57BL/6J dams across the last consecutive breeding cycles. Data are shown as mean ± SD (minimum, maximum) or percentage (n/N), where n indicates the number of animals meeting the specified criterion and N the total number of bred animals. Offspring-related parameters refer to the final litter. Summary litter parameters across all litter per dam are provided in Table S1. PD, postnatal day.

### Metabolic Health

To determine how repeated reproduction affects maternal whole-body energy balance, we measured body weight trajectory and food intake during the final lactational and post-weaning periods; following weaning, energy expenditure and food intake were determined in indirect calorimetry cages, and oral glucose tolerance and insulin sensitivity were assessed; five weeks after weaning, fat and fat-free mass were measured, and brain responsivity to the metabolic hormone leptin was assessed.

#### Four cycles of pregnancy and lactation increased maternal body weight during lactation and during the post-weaning period

Body weight trajectories across the experimental timeline revealed distinct peaks corresponding to successive pregnancies in each parity group (Fig. 1B). One-time pregnant dams exhibited a linear increase in body weight prior to breeding, whereas virgin animals maintained a steady linear trajectory throughout the experiment. Following weaning at postpartum day 21, dams without subsequent litter showed a marked decline in body weight, a pattern not observed in virgin controls.

**Figure 1.**
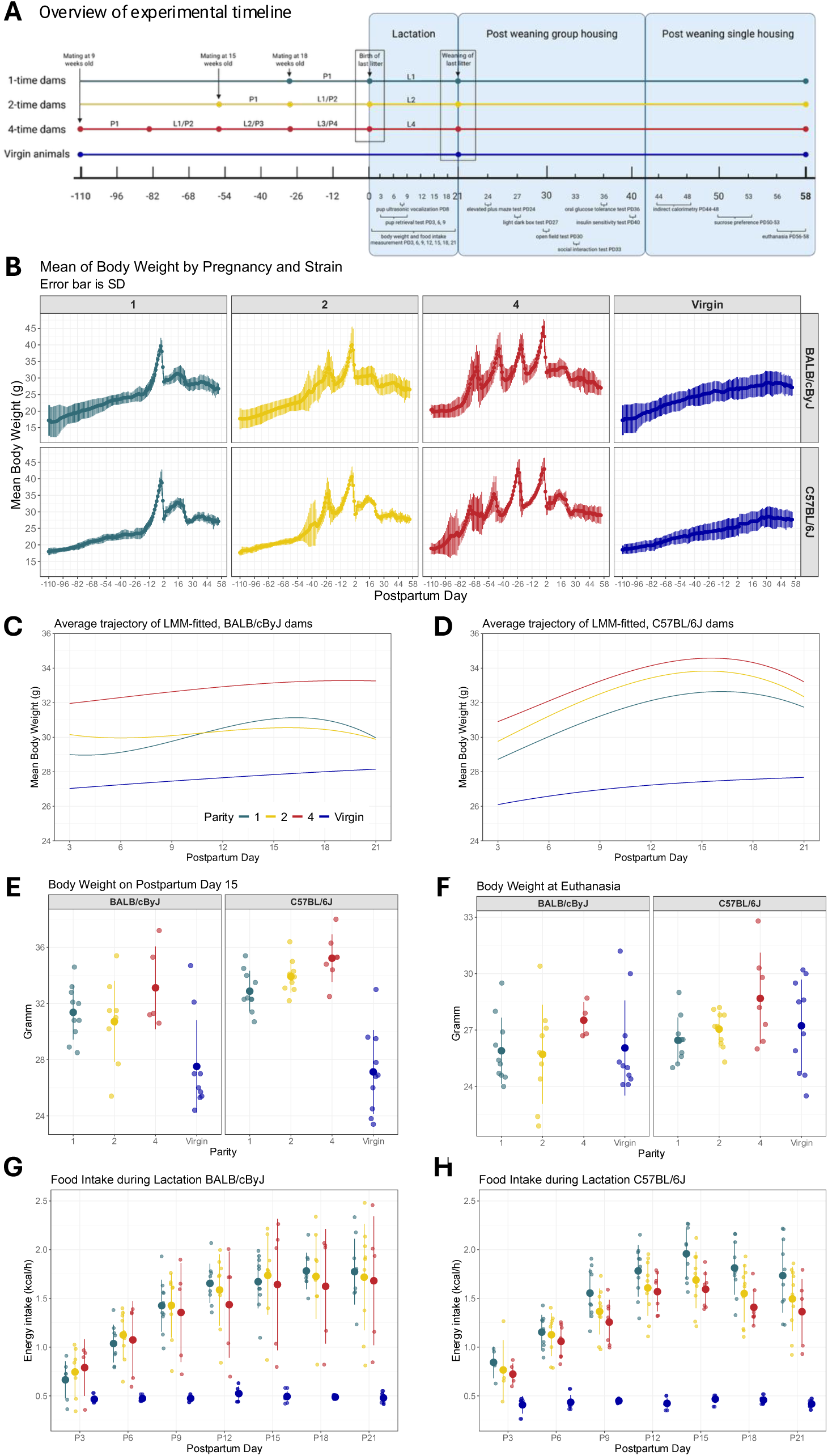
Impact of reproductive load on body weight and food intake across reproductive age and lactation. (A) Overview of experimental timeline in female BALB/cByJ and C57BL/6J dams experiencing one (1-time), two (2-time), four (4-time), or no (Virgin) cycles of pregnancy and lactation. (B) Mean body weight of experimental dams divided by pregnancy level and strain over the whole experimental timeline (postpartum day (PD) -110 to 58). (C, D) Average body weight trajectory during the last lactation as predicted by the linear mixed-effect model (LMM) (SI Appendix, Table S3) in both mouse strains on PD3-21. (E, F) Absolute body weight on PD15 and at termination around PD58. (G, H) Average food intake during the lactation phase in both strains from PD3-21. Data are presented as mean ± SD (B, E-H), or as predictions from LMM (C, D). Dots represent individual animals. Sample sizes for each outcome are provided in SI Appendix, Table S3.

Body weight during the final lactation differed across postpartum days (PD) and varied by both parity and strain (Fig. 1C, D). Consistent with these patterns, main effects of parity number (F(3, 211.71) = 4.04, p = 0.008), strain (F(1, 278.26) = 7.22, p = 0.008), and PD modeled as a third-degree polynomial (PD^3^) (F(2, 426) = 139.83, p < 0.001) were revealed. Interactions between parity × PD^3^ (F(6, 426) = 8.37, p < 0.001), strain × PD^3^ (F(2, 426) = 35.97, p < 0.001), and a three-way interaction between parity × strain × PD^3^ (F(6, 426) = 4.51, p < 0.001) were also found. The number of pups in the last litter did not have an effect on body weight (F(1, 68.95) = 0.50, p = 0.48), nor did the parity × strain interaction (F(3, 272.87) = 0.17, p = 0.92).

Curve morphology was quantified using the cubic coefficient (x^3^), with larger values indicating more pronounced curvature in the body weight trajectory (Table S2). In the BALB/cByJ mice (Fig. 1C), curvature decreased with increasing parity, indicating more dynamic weight gain and loss during the first reproductive cycle and progressively straighter trajectories in subsequent cycles. In contrast, C57BL/6J dams (Fig. 1D) exhibited similar x^3^ values across parity groups, reflecting largely parallel weight trajectories during lactation. Across strains, higher parity was associated with greater initial body weight on PD3. Most dams reached peak body weight on PD15-P16 (Table S2), with the exception of four-time BALB/cByJ dams, which peaked on PD19. Virgin females had near-linear growth and reached their peak body weight on PD21. Because peak body weight during lactation was reached around PD15, body weights were compared at this time point (Fig. 1E). A linear mixed model (LMM) fit at PD15 estimated an average increase of 2.1 g in four-time pregnant dams across both strains compared to one-time pregnant dams, whose estimated weight was 31.4 g (β_4_ = 2.1, 95% CI [0.33, 3.83], p = 0.021; β_1_ = 31.4, 95% CI [30.07, 32.71]). In contrast, virgin animals were on average 4.9 g lighter than one-time pregnant dams (β_Virgin_ = -4.9, 95% CI [-6.37, -3.35], p < 0.001), and C57BL/6J dams were on average 1.6 g heavier than BALB/cByJ dams (β_C57BL/6J_ = 1.6, 95% CI [0.44, 2.73], p = 0.007).

When compared at the terminal timepoint five weeks later (Fig. 1F), females with four reproductive cycles were still on average 1.8 g heavier than one-time pregnant dams (β_4_ = 1.8g, 95% CI [0.53, 3.12], p = 0.007; β_1_ = 25.7g, 95% CI [24.28, 27.04]), and C57BL/6J females were 1.3 g heavier than BALB/cByJ dams (β_C57BL/6J_ = 1.3, 95% CI [0.47, 2.14], p = 0.003). Virgin females, however, were no longer lighter than dams bred one time (β_Virgin_ = 0.33 [-0.76, 1.42] p = 0.549). The elevated body mass in four-time females could not be explained by higher fat mass, as measured by echoMRI after killing (Fig. S1A). Lean mass was increased by 1 gram in two-time pregnant dams compared to one-time pregnant dams, which had an average lean mass of 11.3g, regardless of the strain. C57BL/6J mice were on average 1.1 g heavier than BALB/cByJ animals, which was driven by an average increase in fat mass of 1.1g. Brain sensitivity to the adipokine leptin, as measured by leptin-induced pSTAT3 levels in the hypothalamic arcuate nucleus, was similar across parity and strain five weeks after weaning (Fig. S1B).

#### Food intake increased dramatically during lactation and remained elevated for weeks after weaning, along with energy expenditure

The development of lactational hyperphagia followed a trajectory similar to body weight and peaked around PD15, when dams consumed ∼1.8 kcal/h compared with ***∼***0.5 kcal/h in virgin females, corresponding to an increase of approximately 250% (Figure 1G, 1H). Lactational food intake differed across postpartum days and varied by both parity and strain. Consistent with these patterns and similar to body weight, main effects of parity number (F(3, 324.4) = 45.15, p < 0.001), strain (F(1, 404.9) = 5.12, p = 0.02), and PD modeled as a quadratic (second-degree polynomial) term (PD^2^) (F(2, 379.7) = 722.63, p < 0.001) were revealed. Interactions between parity x PD^2^ (F(6, 380.2) = 100.83, p < 0.001), strain x PD^2^ (F(2, 380.2) = 16.31, p < 0.001), and a three-way interaction between parity x strain x PD^2^ (F(6, 380.2) = 2.19, p = 0.04) were also found. The number of pups in the last litter had a strong influence on maternal food intake (F(1, 65.1) = 219.81, p < 0.001), with each pup increasing maternal food intake by an estimated 0.13 kcal/h (β_pups_ = 0.13 kcal/h, 95% CI [0.11, 0.14], p < 0.001).

Four weeks after weaning, mice were individually housed in indirect calorimetry cages to assess energy intake (Fig. 2A-B), energy expenditure (Fig. 2D-E), and respiratory exchange ratio (RER) (Fig. S1C-D) over three days to investigate whether energy balance remains altered in the post-weaning period. Likelihood-ratio tests detected no significant interactions between body mass and parity, indicating that relationship between body mass and energy expenditure and energy intake were comparable across groups, supporting the use of ANCOVA with body mass as a covariate [22]. Though energy intake remained lower in the virgin females (β_Virgin_ = -0.06, 95% CI [-0.11, -0.01], p = 0.025) than in primiparous females (β_1_ = 0.23, 95% CI [0.04, 0.5]), the difference was markedly smaller than during lactation, and additional pregnancies did not further affect food intake over 24 hours (Fig. 2C). Virgin females also showed lower total energy expenditure over 24 hours (β_Virgin_ = -0.03, 95% CI [-0.05, -0.01], p = 0.003) when compared to one-time pregnant dams (β_1_ = 0.3, 95% CI [0.2, 0.41]), whereas two-, and four-time pregnant dams did not differ from one-time pregnant dams (Fig. 2F). Further, C57BL/6J animals had a lower energy expenditure compared to the BALB/cByJ strain (β_C57BL/6J_ = -0.05, 95% CI [-0.06, -0.03], p < 0.001). Different reproductive experiences did not affect the RER four weeks after weaning, though RER differed by strain during the dark phase, with C57BL/6J mice exhibiting higher RER than BALB/cByJ mice (β_C57BL/6J_ = 0.03, 95% CI [0.01, 0.04], p < 0.001; β_BALB/cByJ_ = 0.96, 95% CI [0.94, 0.98]) (Fig. S1C-D).

**Figure 2.**
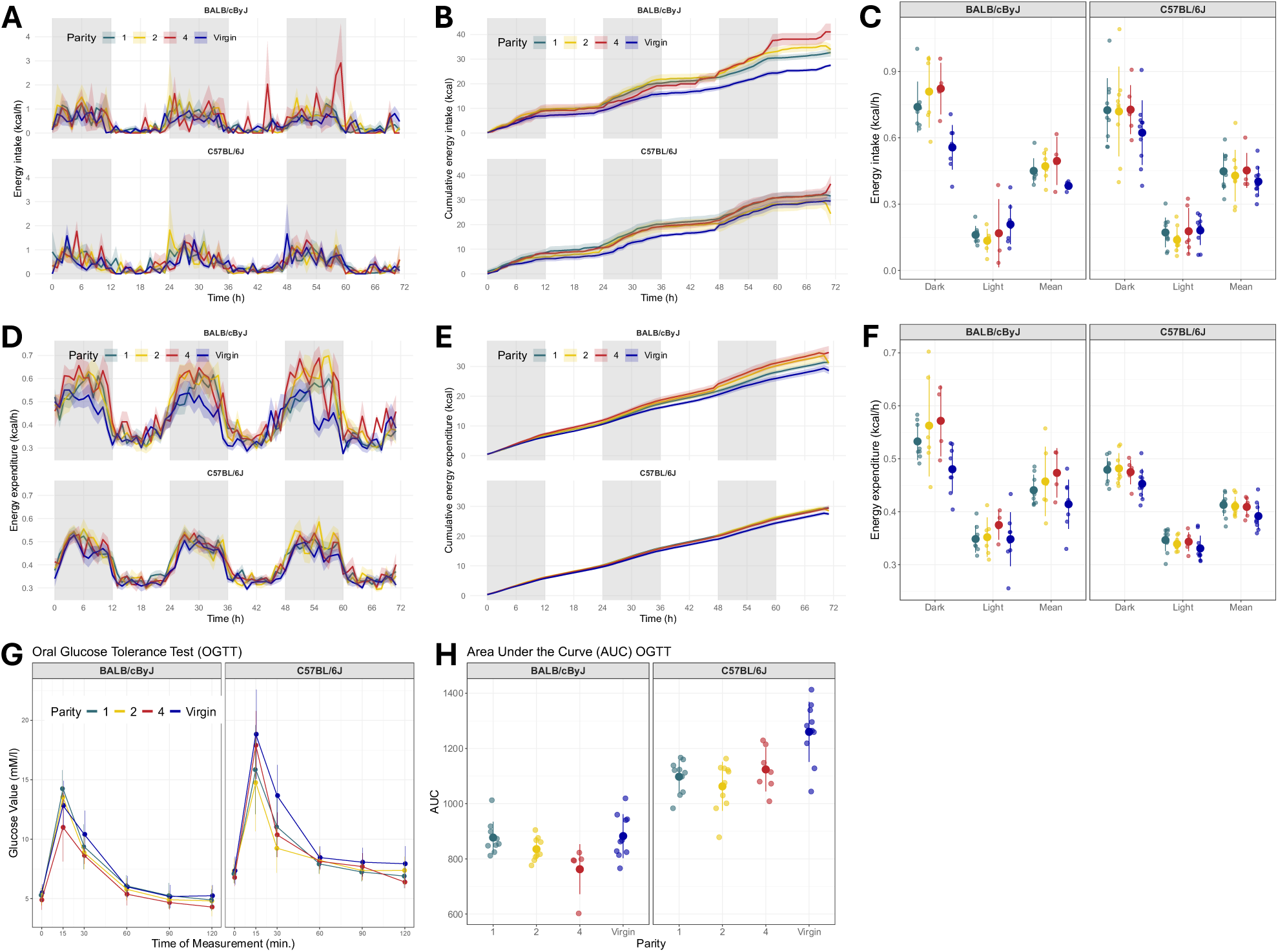
Impact of reproductive load on energy expenditure, food intake and glucose tolerance in the post-lactation period. Female BALB/cByJ and C57BL/6J mice experienced 1, 2, or 4 cycles of pregnancy and lactation, or remained virgin (Virgin). (A, B) Energy intake and cumulative energy intake over three days from postpartum day (PD) 44-48. (C) Mean energy intake (averaged over 3 d) during the dark phase, light phase, and total daily mean. (D, E) Energy expenditure and cumulative energy expenditure over three days from PD44-48. (F) Three-day energy expenditure during the dark phase, light phase, and their combined mean. (G) Blood glucose levels measured around PD 36 at 0, 15, 30, 60, 90, and 120 min following an oral glucose bolus (2.5 g/kg) administered after a 4-hour fast. (H) Area under the curve (AUC) for the glucose tolerance test (OGTT). Data are represented as mean ± SD. Dots represent individual animals (C, F, H). Sample sizes for each outcome are provided in SI Appendix, Table S3.

#### Glucose homeostasis was altered in a strain-specific manner in response to parity numbers

Maternal glucose tolerance and insulin sensitivity were measured three weeks after weaning. C57BL/6J mice exhibited consistently higher blood glucose levels following oral glucose administration than BALB/cByJ mice (Fig. 2G), resulting in elevated AUC values (Fig. 2H) (β_C57BL/6J_ = 220.7, 95% CI [151.3, 290.2], p < 0.001; β_BALB/cByJ_ = 878.0, 95% CI [828.1, 927.9]). Further, there was evidence of interaction between parity and strain, suggesting strain-specific adaptations in glucose clearance with increasing parity cycles. Four-time pregnant BALB/cByJ animals had decreased AUC values compared to one-time BALBc/ByJ (β_4_ = -116.1, 95% Cl [-201.2, -31.1], p = 0.008), whereas virgin animals had similar AUC values compared to one-time BALBc/ByJ dams (β_Virgin x BALBc/ByJ_ = 4.5, 95% Cl [-64.7, 73.7], p = 0.9). In the C57BL/6J strain, AUC values were comparable across all parity groups, with an increase in virgin animals (β_Virgin x C57BL/6J_ = 157.1, 95% Cl [59.1, 255.0], p = 0.002). Insulin sensitivity (Fig. S1E-F) was not altered by parity, but as observed during the OGTT, C57BL/6J generally had higher plasma glucose values than BALB/cByJ animals.

### Maternal and exploratory behavior

To assess the impact of reproductive load on maternal behavior, exploratory behavior, and locomotor activity, dams underwent a battery of behavioral assays. The following tests were conducted: Five minutes pup retrieval on PD3, PD6, PD9; five minutes elevated plus maze test (EPM), ten minutes light-dark box (LDB) paradigm, thirty minutes open field test (OF), ten minutes social interaction test (SI), 23 hours sucrose preference test (SP), and histological examination of oxytocin positive neurons in the paraventricular nucleus of the hypothalamus (PVN).

#### Maternal motivation, assessed by pup retrieval during lactation, was strain-dependent but unaffected by parity

A subset of BALB/cByJ dams failed to retrieve the first pup within the test period on PD9 (3 of 69 tests), whereas all C57BL/6J dams retrieved the first pup across all postpartum days (78 of 78 tests) (Fig. 3A). Additionally, 20% of BALB/cByJ dams did not retrieve all displaced pups to the nest (14 of 69 tests), compared with 3% of C57BL/6J dams (2 of 78 tests) (Fig. 3B). C57BL/6J mice exhibited shorter latencies than BALB/cByJ for both first pup retrieval (36% reduction, 95% CI [19–50%], p < 0.001; β_C57BL/6J_ = −0.45) and total retrieval duration (31% reduction, 95% CI [11–47%], p = 0.004; β_C57BL/6J_ = −0.37). Latency decreased across postnatal days, with significant reductions at PD9 for first pup retrieval (28% reduction, 95% CI [13–40%], p < 0.001; β_PD9_ = −0.32) and at both PD6 (20% reduction, 95% CI [7–31%], p = 0.005; β_PD6_ = −0.22) and PD9 (35% reduction, 95% CI [24–44%], p < 0.001; β_PD9_ = −0.42) for total retrieval duration. Parity had no effect on either measure. Consistent with these findings, total nest time was almost twice as long in the C57BL/6J strain compared to the BALB/cByJ strain (β_C57BL/6J_ = 65.9 s, 95% CI [49.0, 82.9], p < 0.01; β_BALB/cByJ_ = 79.4 s, 95% CI [58.7, 100.1]) (Fig. 3C). Further, there was a tendency of slightly increased nest attendance of 20.4 s in four-time pregnant dams compared to one-time pregnant dams across both strains (β_4_ = 20.4 s, 95% CI [-1.5, 42.3], p = 0.07; β_1_ = 79.4 s, 95% CI [58.7, 100.1]), as well as a tendency for shorter nest attendance on PD6 compared to PD3 (β_PD6_ = -20.3 s, 95% CI [-41.2, 0.6], p = 0.06; β_PD3_ = 79.4 s, 95% CI [58.7, 100.1]). No effect of pup sex on first retrieval was observed (data not shown).

**Figure 3.**
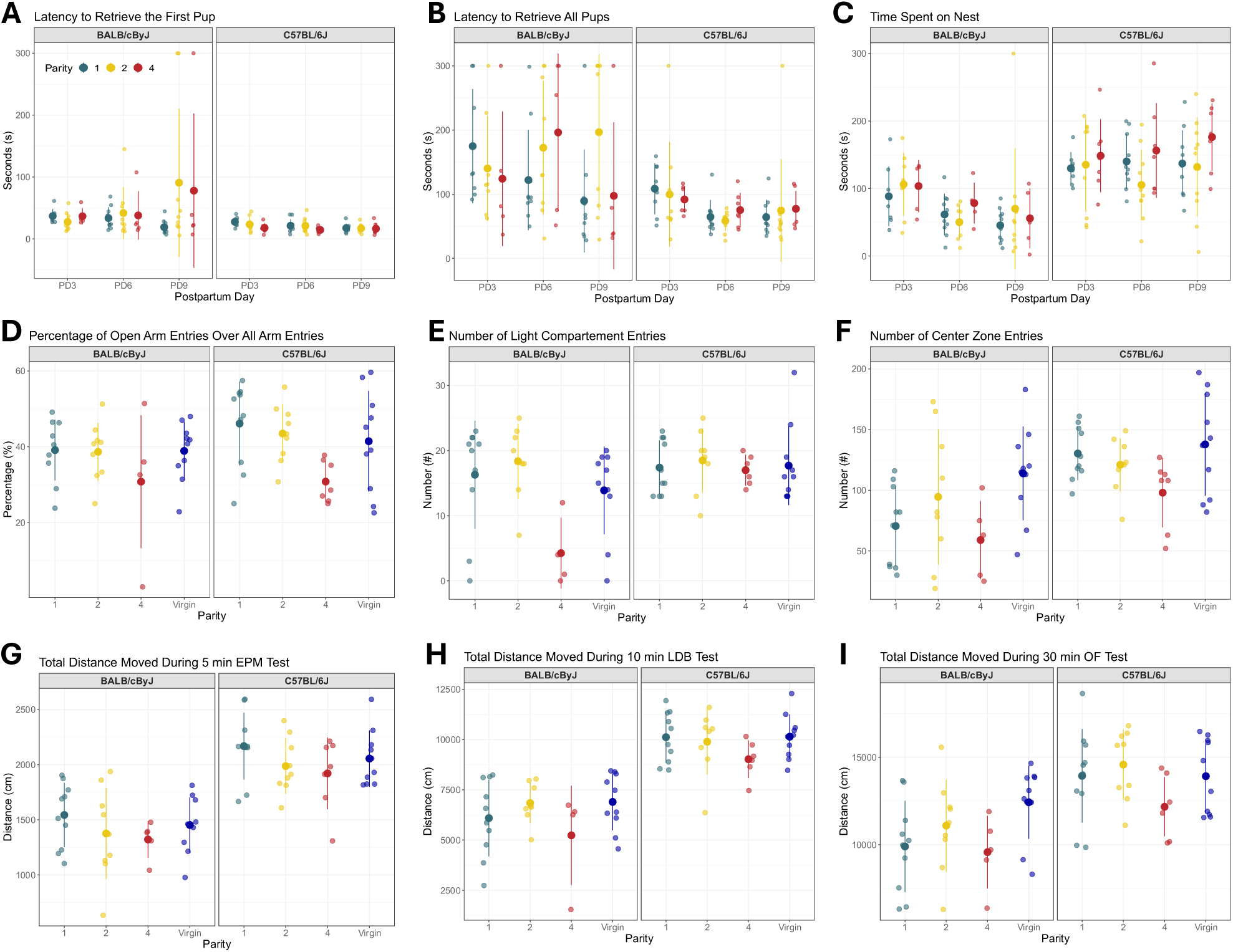
Impact of reproductive load on maternal behavior, exploratory behavior, and locomotor activity in the post-lactation period in female BALB/cByJ and C57BL/6J dams experiencing 1, 2, 4, or no (Virgin) cycles of pregnancy and lactation. Latency to retrieve the first pup (A), all the pups (B), and time spent on the nest (C) during a five-minute pup retrieval test on postpartum day (PD) 3, 6, and 9. Percentage of open arm entries (D) and total distance moved (G) during a 5-minute elevated plus maze test (EPM) conducted around PD24. Number of light-compartment entries (E) and distance moved (H) during a 10-minute light-dark-box (LDB) test paradigm run around PD27. Number of center zone entries (F) and distance moved (I) during a 30-minute open field (OF) test around PD30. Data are represented as mean ± SD. Dots represent individual animals. Sample sizes for each outcome are provided in SI Appendix, Table S3.

#### Multiparity was associated with reduced exploratory behavior

Across three behavioral tests, four-time dams consistently exhibited a reduction in both exploratory behavior and activity compared to primiparous dams. Four-time dams entered the open arms of the EPM 12.1% fewer times on average (β_4_ = -12.1%, 95% CI [-19.3, -5.0], p = 0.001; β_1_ = 40.6%, 95% CI [35.7, 45.6]) (Fig. 3D), and correspondingly spent 12.8% less time in the open arms (β_4_ = -12.8%, 95% CI [ -23.2, -2.5], p = 0.016; β_1_ = 36.4%, 95% CI [29.2, 43.6])( Fig. S2A). Further, four-time pregnant dams had an overall reduction in the total distance travelled during the EPM test, with a tendency for reduction in two-time pregnant dams when compared to one-time pregnant dams (β_4_ = -235.9 cm, 95% CI [-445.8, -25.9], p = 0.028; β_2_ = -174.7 cm, 95% CI [-361.2, 11.7], p = 0.07; β_1_ = 1550.9 cm, 95% CI [1405.2, 1696.5]) (Fig. 3G). During the LD test, BALB/cByJ four-time pregnant dams entered the light compartment 12.1 times less compared to one-cycle dams (β_4 x BALB/cByJ_ = -12.1, 95% CI [ -19.1, -5.1], p = 0.001; β_1 x BALB/cByJ_ = 16.3, 95% CI [12.6, 20.1]) (Fig. 3E) and spent less time in the light box (Fig. S2B). Four-time-pregnant dams also tended to show a reduction in total distance moved (β4 = -949.9 cm, 95% CI [ -2070.1, 170.2], p = 0.095) (Fig. 3H). In the OFT, four-time dams also showed a 27% reduction in the number of center zone entries (β_4_ = -21.8%, 95% CI [-48.2, 4.7], p = 0.105) (Fig. 3F) and in time spent in the center zone (Fig. S2C), though the reduction in total distance moved was not as pronounced as in the earlier tests when compared to one-time dams (β_4_ = -1074.5cm, 95% CI [-2762.3, 613.4], p = 0.208; β_1_ = 10457.9 cm, 95% CI [9287.0, 11628.7]). (Fig. 3I)

A strong strain effect was also observed across all tests, with C57BL/6J mice showing greater locomotor activity than BALB/cByJ mice. In the EPM (Fig. 3G), C57BL/6J mice travelled 39% more distance than BALB/cByJ mice (β_C57BL/6J_ = 611.4 cm, 95% CI [473.8, 748.9], p < 0.001; β_BALB/cByJ_ = 1550.9 cm, 95% CI [1405.2, 1696.5]). Similar patterns were observed for the LD and OFT (Fig. 3H and 3I), where C57BL/6J mice moved 50% and 28% greater distances, respectively, than BALB/cByJ C57BL/6J dams entered the center zone 46% more often compared to the BALB/cByJ strain (β_C57BL/6J_ = 37.4, 95% CI [20.1, 54.7], p < 0.001; β_BALB/cByJ_ = 81.7, 95% CI [63.3, 100.1]) (Fig. 3F).

Social interaction was not affected by the number of pregnancy and lactation (Fig. S2D-F). However, a strong strain effect was observed, with C57BL/6J dams spending more time interacting with both the prisoner and the dummy mouse and moving overall less than BALB/cByJ dams. Anhedonia-like behavior, assessed by a sucrose preference test around 30 days post-weaning, was unaffected by reproductive experience but differed by strain (Fig. S2G). C57BL/6J animals had, on average, a 16.8% higher sucrose preference compared to the BALB/cByJ strain. Oxytocin-positive cell counts in the PVN varied with reproductive history five weeks after weaning the last litter. Both four-time dams and virgin animals exhibited fewer oxytocin-positive cells compared to one-time dams across both strains (Fig. S2H).

### Nutritional and hormonal status

We next quantified physiological markers of maternal nutritional and hormonal status, including fecal corticosterone metabolites, skeletal parameters, liver iron concentration, and gene expression profiles in liver and adipose tissue.

#### Fecal corticosterone metabolites (FCMs) were dependent on strain and sampling time points but were not affected by reproductive history

FCMs were quantified at baseline, weaning, and killing to assess HPA axis activity (Fig. 4A). Reproductive experience did not influence FCM levels at any time point, suggesting that the number of breeding cycles did not alter basal or dynamic HPA axis activity. In contrast, strong strain differences were observed across all time points, with C57BL/6J females having approximately 65% lower values relative to BALB/cByJ (ratio = 0.35; log-scale β_C57BL/6J_ = -1.0, 95% CI [-1.4, -0.7], p < 0.001). The FCM trajectories also differed by strain. While BALB/cByJ animals showed a trend toward reduced FCM levels at the weaning time point compared to baseline (log-scale β_BALB/cByJ x weaning_ = -0.3, 95% CI [-0.6, 0.0], p = 0.067; log-scale β_BALB/cByJ x baseline_ = 5.4 CI [5.1, 5.6]), C57BL/6J animals exhibited significantly increased levels at weaning (log-scale β_C57BL/6J x weaning_ = 1.0, 95% CI [0.6, 1.5], p < 0.001). Absolute FCM concentrations (ng/50 mg feces) are provided in Fig. S3A. Baseline levels, measured before breeding pair setup, were not associated with first- or multiparous breeding success rates (data not shown).

**Figure 4.**
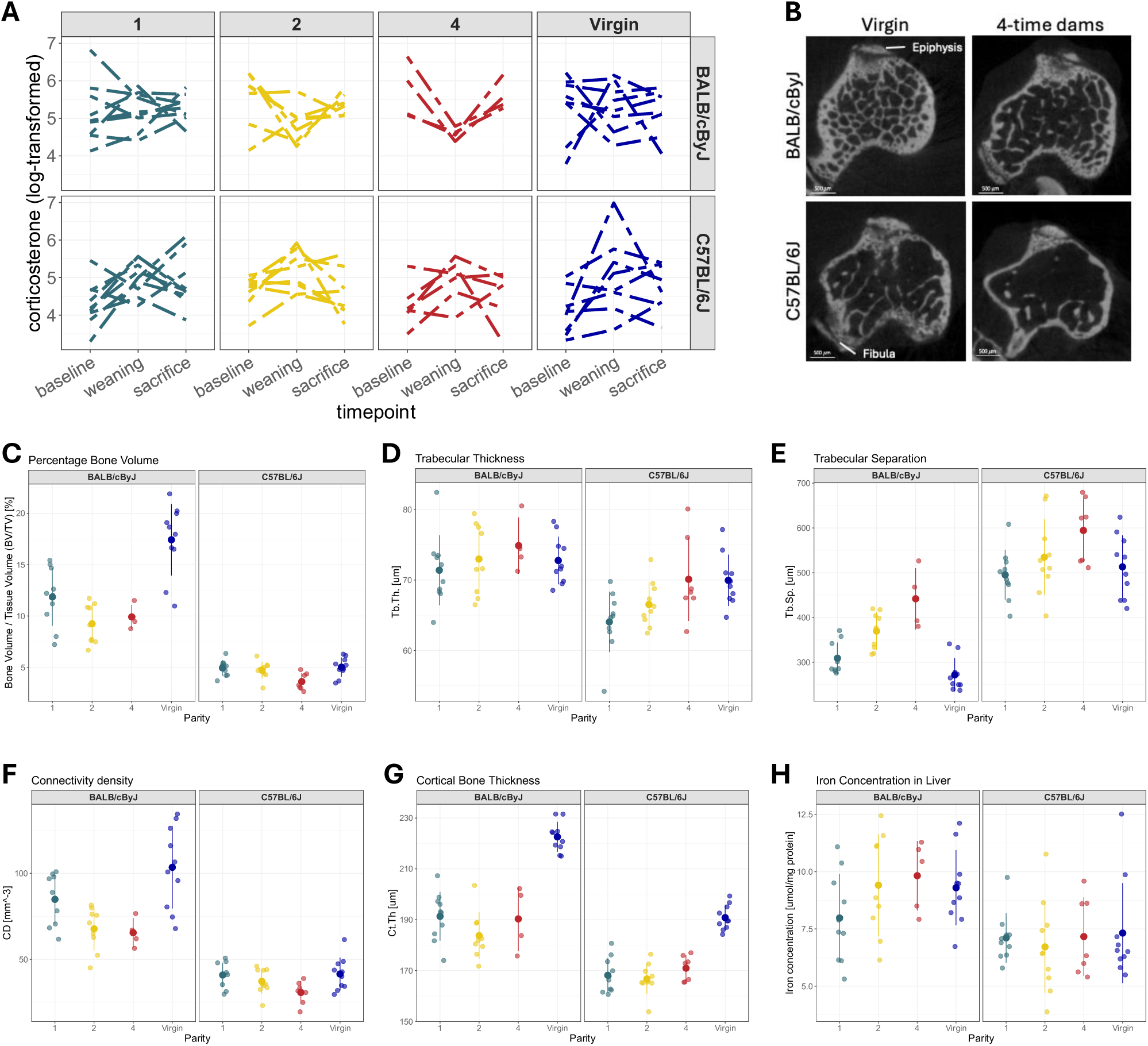
Impact of reproductive load on fecal corticosterone metabolites and bone structure in female BALB/cByJ and C57BL/6J animals experiencing 1, 2, 4, or no (Virgin) cycles of concurrent pregnancy and lactation. (A) Log-transformed fecal corticosterone metabolites (FCMs) levels before breeding onset (baseline), at weaning of the last litter (weaning), and on the termination day of the animals (sacrifice). (B) Representative images of microcomputer tomography (micro-CT) scans of the proximal tibia in 4-time reproducing dams and virgin control animals across both strains. Bone volume fraction (BV/TV) (C), trabecular thickness (Tb.Th) (D), trabecular separation (Tb.SP) (E), connectivity density (CD) (F), and cortical bone thickness (Ct.Th) (G) of reproducing dams and virgin animals after termination around postpartum day (PD) 58. (H) Liver iron concentration at termination around PD58. Data are presented as lines representing individual animals measured longitudinally across three time points (A) or as mean ± SD with dots representing individual animals (C-H). Sample sizes for each outcome are provided in SI Appendix, Table S3.

#### Bone mass differed between strains and showed a dose-dependent response to reproductive experience, with progressive changes across increasing parity levels

Micro-CT analysis of proximal tibiae collectively revealed a progressive weakening of bone structure with increasing reproductive cycles, characterized by reduced bone volume, loss of the bone trabecular architecture, and concomitant decreased connectivity density (Fig. 4C-4F). An interaction between strain and parity indicates a strain-specific response to pregnancy and lactation. In BALB/cByJ mice, two cycles of reproduction decreased bone volume by 2.6% compared to one cycle (β_BALB/cByJ x 2_ = -2.6%, 95% CI [ -4.4, -0.8], p = 0.005), while bone volume was 5.6% higher in virgin females (β_BALB/cByJ x Virgin_ = 5.6%, 95% CI [3.8, 7.3], p < 0.001) (Fig. 4C). These effects were not observed in C57BL/6J females, who exhibited 6.9% lower bone volume fraction than BALB/cByJ females (β_C57BL/6J_ = -6.9%, 95% CI [-8.6, -5.2], p < 0.001; β_BALB/cByJ_ = 11.9%, 95% CI [10.6, 13.1]). Across both strains, trabecular thickness and trabecular separation (Fig. 4D-E) increased with parity, consistent with progressive changes in trabecular architecture. Trabecular number showed a similar strain- and parity-dependent pattern, decreasing in BALB/cByJ dams with two and four cycles and in C57BL/6J dams, while increasing in virgin BALB/cByJ mice (Fig. S3B). Connectivity density, describing how well trabeculae are interconnected within the trabecular bone network, showed pronounced strain specificity, with C57BL/6J dams exhibiting ∼50% lower values than BALB/cByJ animals (_C57BL/6J_ = -44.0 mm^-3^, 95% CI [-55.6, -32.4], p < 0.001; β_BALB/cByJ_ = 84.9 mm^-3^, 95% CI [76.7, 93.1]) (Fig. 4F). In BALB/cByJ mice, connectivity decreased in two- and four-time dams and increased in virgins relative to one-time dams (β_4 x BALB/cByJ_ = -19.4 mm^-3^, 95% CI [-34.8, -4.0], p = 0.014; β_2 x BALB/cByJ_ = -17.1 mm^-3^, 95% CI [-29.1, -5.2], p = 0.006), whereas no parity effects were detected in C57BL/6J mice. Cortical thickness was reduced in dams compared to virgins across both strains (Fig. 4G), with an additional strain-specific reduction in two-time pregnant BALB/cByJ mice (β_2_ = -7.6 μm, 95% CI [-14.4, -0.8], p = 0.029; β_1_ = 191.3 μm, 95% CI [186.6, 196.0]). Overall, C57BL/6J mice exhibited lower cortical thickness than BALB/cByJ mice (β_C57BL/6J_ = -23.2 μm, 95% CI [-29.8, -16.6], p < 0.001). Representative micro-CT images illustrating these strain- and parity-dependent differences are shown in Fig. 4B, highlighting consistently higher bone mineral density in BALB/cByJ and virgin mice, compared to C57BL/6J and multiparous mice. Given the differences in body weight observed at termination, tibia length was assessed to determine whether overall skeletal growth differed between groups. Tibia length did not differ between parity groups or strain (Fig. S3C).

#### Hepatic iron levels, measured as an indicator of systemic iron stores five weeks after weaning, exhibited no effect of reproductive history

C57BL/6J females showed consistently lower liver iron concentrations compared to BALB/cByJ females (b_C57BL/6J_ = -2.0 mmol/mg, 95% CI [-2.8, -1.1], p < 0.001; b_BALB/cByJ_ = 8.5 mmol/mg, 95% CI [7.6, 9.5]) (Fig. 4H). In contrast, at the gene expression level, iron metabolism appeared to be regulated differently between reproductive and non-reproductive states, with virgin animals expressing higher levels of TRF1, FPN1, IRP1, and GPX4 compared to reproducing dams (Fig. S3D). Hepatic genes related to fatty acid (FASN, SCD1, Ppara), cholesterol (ApoB, Cyp7a1), and glucose metabolism (G6Pase, FGF21) were often downregulated in multiparous dams (Fig. S3E-F). In adipose tissue, fatty acid metabolism genes (FASN, SCD1, Steap4, Ppara) and adipokines (Leptin, FGF21, Fabp4, Resistin) showed minimal differences, except for an upregulation of FGF21 and Resistin in multiparous dams (Fig. S3G-H).

## Discussion

Despite the widespread use of rodent breeders to support biomedical research, how female breeders adapt in response to repeated reproductive efforts has been poorly characterized. Here we report findings from an exploratory study designed to capture a composite picture of maternal health spanning reproductive, immediate-post-reproductive, and recovery phases in two commonly used inbred strains. Some readouts showed clear and dose-dependent responses to increasing reproductive load. Others were modified by a single cycle of reproduction and did not change further. Still others remained stable across parity levels. Strain differences were pervasive, underscoring that reproductive adaptation is not uniform even across standard laboratory models. These findings are discussed in the context of maternal resilience and its limits under intensive breeding conditions.

A notable feature of the study design warrants consideration before interpreting the parity-dependent findings. Although dams were randomly assigned to reproductive conditions, the proportion successfully completing the assigned number of cycles varied considerably by group and strain, with only 14% of BALB/cByJ and 23% of C57BL/6J dams reaching the four-cycle endpoint. Our inclusion criteria required dams to complete each assigned cycle to remain in the study, meaning that the four-cycle group represents a self-selected subset of highly reproductively capable females, and that the costs of intensive breeding reported here may therefore be conservative estimates. The question of what distinguishes dams that can sustain four cycles from those that cannot remain open and worthy of investigation, and this natural variation in reproductive capacity may itself reflect meaningful biological diversity [23]. It is also worth noting that the study was conducted during a period of significant facility construction, which introduced chronic background noise and vibration. While this represents an uncontrolled variable, it also reflects real-world conditions in active research facilities. The fact that dams reproduced successfully under these circumstances speaks further to their resilience.

### Reproductive load reshapes maternal energy balance and glucose tolerance, but with little evidence of metabolic dysfunction

In women, lactation is supported largely by fat stores accumulated during pregnancy, such that postpartum body mass tends to decline across the lactation period [24,25]. Unlike in humans, mouse dams have minimal capacity to buffer the energetic demands of lactation through fat mobilization, instead meeting an approximate three- to four-fold increase in energy demand through hyperphagia and substantial organ remodeling, including the gut [3,9,26]. The peak in body mass observed around PD15-16 in both strains is therefore best understood as the point of maximal metabolic and structural adaptation to lactation, rather than a sign of excess energy storage. The two strains responded differently to increasing reproductive load: BALB/cByJ dams reach peak body mass later (PD19) with higher parity, while C57Bl/6J dams show a progressively higher peak with each reproductive cycle. Notably, the elevated body mass persisting in four-cycle dams until the end of the study could not be attributed to increased fat mass. This raises the intriguing possibility that the capacity for progressive organ remodeling may not only be strain-specific but might also be a feature of the reproductively resilient females—that those able to continue adapting metabolically are precisely those capable of sustaining the highest reproductive demand.

As expected [26,27], all dams showed marked increases in food intake during lactation, but multiparous dams consumed less food than primiparous dams despite their greater body mass. This inverse relationship suggests that factors beyond food intake drive lactational weight gain in experienced breeders. Litter size was also a strong independent predictor of food intake, reflecting the known relationship between nursing demand and the magnitude of maternal hyperphagia [3,26,27]. In the weeks following weaning, parous females of both strains maintained higher food intake and energy expenditure than virgin controls, but neither was further modified by the number of reproductive cycles. These findings suggest that a single cycle of reproduction is sufficient to shift maternal energy turnover to a new baseline, and that shift is not compounded by additional rounds of breeding.

Three weeks after weaning, C57BL/6J dams exhibited consistently higher blood glucose and greater AUC following oral glucose challenge than BALB/cByJ dams, extending to female mice the established strain difference in glucose metabolism previously described in males [28–30]. Parity also influenced glucose clearance in a strain-specific manner. Four-cycle BALB/cByJ dams showed improved glucose clearance relative to primiparous dams, while virgin animals were indistinguishable from primiparous dams. In C57BL/6J dams, AUC values were comparable across parity groups, with virgin animals showing elevated AUC relative to primiparous dams, suggesting that even a single cycle may improve glucose handling in this strain. Together, these data suggest that reproductive experience can drive improvements in glucose handling. This is consistent with evidence in women showing that breastfeeding reduces long-term risk of type 2 diabetes [31], and with one study in mice showing that two spaced rounds of pregnancy improved glucose tolerance up to three months after weaning [32].

### Mouse dams exhibit behavioral adaptations to increasing reproductive load, though maternal responsiveness and stress hormone levels are not influenced by parity

Pup retrieval, a readout of maternal responsiveness known to be sensitive to social and environmental stressors [12,13], was unaffected by reproductive experience. These findings suggest that even under high reproductive demand, dams prioritize and maintain the behaviors most critical to offspring survival. Strain differences were again evident, with C57BL/6J dams retrieving faster and more consistently than BALB/cByJ dams, as previously observed [33].

In contrast, exploratory and locomotor activity were progressively reduced in multiparous dams across three behavioral assays, with the strongest effect observed in the elevated plus maze conducted closest to weaning. This reduction could reflect a cumulative shift in exploratory drive because of high reproductive load, or an adaptive tendency toward energy conservation in dams with more reproductive experience. Given the self-selected nature of the four-cycle group, however, it is also possible that a more conservative behavioral phenotype is a pre-existing trait in females best equipped to sustain repeated reproduction, rather than a consequence of it. Consistent with observations in female mice used repeatedly for embryo transfer [34], FCM levels were not influenced by reproductive history, only by strain and sampling time point. These findings argue against chronic physiological stress as an explanation for reduced exploratory behavior, and instead support the interpretation that reduced exploration reflects an adaptive rather than maladaptive phenotype.

### Reproductive experience progressively remodels bone microstructure, while hepatic iron levels appear restored after weaning

Skeletal tissue showed a clear dose-dependent response to reproductive load. Analysis of the proximal tibiae revealed progressive changes in trabecular architecture across parity levels in both strains, including reduced bone volume, increased trabecular separation, and decreased connectivity density, with effects detectable from the first reproductive cycle onward. While these findings align with previous data collected from dams with spaced pregnancies [19], the enhanced cortical thickness observed in that study was not replicated here, raising the question of whether consecutive breeding cycles may compromise load-bearing capacity in ways that spaced pregnancies do not.

In contrast, hepatic iron concentration was unaffected by reproductive history five weeks after weaning, suggesting that iron stores are restored within this timeframe. Exploratory analysis of iron metabolism gene expression, however, revealed higher expression of several iron regulatory genes in virgin compared to parous animals across both strains, which presents the possibility that repeated reproduction leaves a lasting impact on iron handling that persists beyond the restoration of iron levels.

### Breeding shapes the mouse dam—weighing costs and potential benefits

Taken together, the data paints a complex picture of life as a breeding mouse dam. Reproductive experience leaves a measurable imprint on maternal physiology, most clearly in bone microstructure, and more subtly in body composition, energy balance, and glucose homeostasis. Readouts of maternal motivation, however, remained largely stable across levels of reproductive load, and the absence of elevated fecal corticosterone metabolites in multiparous dams argues against chronic physiological stress as a feature of the intensive consecutive breeding studied here. The self-selected nature of the four-cycle group complicates simple interpretation of these costs: while the reproductively resilient females may underestimate the average physiological impact, those that did not complete four cycles may have been protected by an adaptive tendency to limit reproductive output. Still, reducing the experience of breeding to its physiological costs alone would be incomplete. Maternal behavior is one of the most strongly motivated behaviors in mammals, engaging powerful brain reward circuitry [35,36], and the opportunity to express this behavior represents a form of enrichment that perhaps has not been fully recognized in the context of laboratory breeding. Emerging discussions around positive animal welfare [37,38] are beginning to provide the framework to recognize this.

These findings suggest that good welfare for the breeding dam encompasses more than the absence of suffering, and that both the physiological and behavioral components of her experience deserve consideration. Strain differences were present in nearly every readout, highlighting just how different two inbred strains of mice can be, even in response to something as natural as reproduction, which cautions against generalizing findings from a single model or strain [39,40]. Notably, the comprehensive characterization of virgin females alongside breeding dams in both strains contributes a useful baseline dataset for preclinical research in females. Together, these observations push us to consider that optimal breeding conditions may not be uniform across strains, and that more conscious and strain-specific breeding strategies could benefit both the dam and broader breeding goals like reducing surplus animals. We hope these data contribute to a more deliberate consideration of breeding dam welfare—not just as a peripheral concern of the research they support, but as a subject of scientific inquiry that deserves attention in its own right.

## Methods

### Animal Subjects and Housing

Experimental mice were derived from founding breeders of the inbred strains BALB/cByJ and C57BL/6J (Charles River, Germany) and maintained in-house for at least one generation. The study was conducted in five sequential batches to improve reproducibility [41] and to allow first-litter offspring to serve as breeders for subsequent batches, thereby reducing surplus animals. Animals were randomly allocated (randomizer.org) to one of four experimental groups: one cycle of pregnancy and lactation [1-time dams], two consecutive cycles [2-time dams], four consecutive cycles [4-time dams], and virgins [Virgin]. All pups were weaned on postpartum day 21 (PD21), corresponding to a three-week lactation period. Mice were housed in individually ventilated cages (IVC type II long cages, 365 x 207 x 140mm) with wood-shaving bedding, nesting material, enrichment (crinkle paper, wooden gnawing sticks), and red plastic houses. Mice had ad libitum access to water and standard breeding chow (3.415 kcal/g; Breeding Extrudate 3336; Kliba Nafag; Switzerland). Animals were group-housed and bred in a temperature and humidity-controlled facility (21 ± 1°C, 55 ± 5%) under a reversed 12-h light–dark cycle (lights off: 11 am – 11 pm) and were handled using non-aversive cup handling whenever possible. All procedures were approved by the Cantonal Veterinary Office Zurich, Switzerland (license numbers ZH087/2019 and ZH022/2023) and reported in accordance with the ARRIVE guidelines[42].

### Breeding

Approximately five days prior to mating, baseline feces (2 - 8 pellets), urine (150-200 μl), and blood samples (200 μl or max. 1% of body weight) were collected. Fresh feces and urine were collected by allowing the mouse to defecate and urinate spontaneously into a plastic beaker or onto plastic wrap. Samples were transferred to 0.5 ml Eppendorf tubes and kept on ice. Blood was collected from either the lateral or medial saphenous vein using a 20-gauge needle to puncture the blood vessel, transferred to a Li-heparin tube (Sarsted, Nümbrecht, Germany), and kept on ice until centrifugation for 10 minutes at 10’000 rpm (4°C). Plasma was collected and, together with fecal and urine samples, stored at -80°C until further processing.

Female mice were subjected to a timed mating procedure defined by their parity group. Breeding groups were established by first adding soiled male bedding to the females’ cage two days prior to mating to induce estrus (Whitten effect [43]). The pairing of non-sibling breeders was staggered to align the different cycles of pregnancy and lactation, ensuring age-matching at the study endpoint. Virgins and 4-time dams were paired at 9 weeks of age with either another female mouse (Virgins) or a male mating partner from a different litter (4-time dams) (Fig. 1A). Two-time dams were mated at 15 weeks, and 1-time dams at 18 weeks of age. All animals were housed in same-sex group cages until mating. As soon as breeding pairs were housed together, body, food, and litter weight (if litter present) were collected three times per week, regardless of postpartum day of the dam (Monday, Wednesday, Friday). The presence of pups was checked daily starting from the anticipated gestation day (GD) 17 and designated as postpartum day 1 (PD1) upon detection of pups. On PD1, the number of live and dead pups was noted with as little disturbance of the dam and pups as possible. On PD2 and PD21, dam’s body weight, litter size and weight, pup sex, and total male and female pup weight were noted. Two days before the expected birth of the last litter, male mating partners were removed to prevent subsequent pregnancy during postpartum estrus. During the final lactation, body weight (g), food intake (kcal/h), and litter weight (g) were collected on: PD2, PD3, PD6, PD9, PD12, PD15, PD18, and PD21. Final group sizes (BALB/cByJ, C57BL/6J) were as follows: parity 1 (10,10), parity 2 (9,10), parity 4 (5,7), and virgins (10,10). One BALB/cByJ dam in the four-cycle group was excluded from subsequent analysis after behavioral assessment due to reproductive pathology (vaginitis). At weaning of the final litter on PD21, feces and urine samples were collected as described previously and stored at -80°C until further processing.

### In vivo procedures during the final reproductive cycle

During the final lactation (4-time, 2-time, 1-time dams), pup retrieval tests lasting for five minutes were conducted on PD3, PD6, and PD9, followed by home-cage behavior recordings on PD3, PD6, PD9, PD12, PD15, and PD18 lasting for one hour in both the dark and light phases (2 pm and 2 am). Home-cage behavior recordings were also collected of the paired Virgin females at equivalent intervals. On PD8, an isolation-induced pup ultrasonic vocalization (USV) test was conducted on one male and one female pup per litter, if available. Analysis of home-cage recordings and USV are not included in this manuscript. After final weaning of pups on PD21, dams were housed in mixed-condition groups of 2-5 mice, matched by postpartum day (maximum 7 days apart), for further testing. From this time point onwards, experimenters were blinded to the final reproductive cycle number of dams. For all behavioral and metabolic assessments, tests were ordered based on increasing disruptiveness (i.e. from the minimally disruptive EPM to the maximally disruptive single housing for indirect calorimetry) with at least three days between tests. On an individual testing day, subject order was semi-random, with consideration given to alternating strains.

Following weaning, behavioral phenotyping tests were performed on the following days (± 3 days) to assess the behavioral consequences of multiple rounds of breeding: Elevated plus maze (EPM) test on P24, light-dark-box (LD) test on P27, open field test (OFT) on P30, and a social interaction (SI) test on P33. Behavioral testing was followed by metabolic assays (± 4 days). An oral glucose tolerance test (OGTT) was performed on P36, followed by an insulin sensitivity test (IST) on P40. On P42, mice were housed singly in an indirect calorimetry system (PhenoMaster; TSE Systems, Bad Homburg, Germany) to measure the respiratory exchange ratio (RER), energy expenditure (EE), and daily food intake for three consecutive days. To conclude in vivo experiments, a sucrose preference test (SPT) was conducted from days P51 to P53. Dams were then reintroduced to group housing before study end on P55 (± 5 days). Detailed protocols for all behavioral and metabolic assays are provided in the Supplemental Materials.

### Tissue collection and processing

On the last experimental day, dams were weighed, and fecal and urine samples were collected and stored at -80°C as described previously. After a two-hour fast during the light cycle, dams were reweighed and given an intraperitoneal injection of leptin (3 mg/kg, PeproTech, Cat. #450-31-5MG, Lot # 012176 A2821). Forty minutes later, mice were killed by pentobarbital overdose (300-450 mg/kg, intraperitoneal, Formula magistralis, 50 mg/ml, Cantonal Pharmacy of Zurich, Switzerland). Terminal blood was sampled using cardiac puncture, centrifuged (10 min at 10’000 rpm at 4°C), and plasma was frozen at -80°C. Ear and liver tissue from the left and right lateral lobes were collected and flash frozen in liquid nitrogen. Prior to perfusion, the aorta abdominalis was clamped to prevent perfusion of the lower body. Mice were flushed by peristaltic pump (flow rate 20 rpm) via the left ventricle with 0.1 M ice-cold phosphate buffer (PB, pH 7.4) for 1.5 minutes, followed by a 2.5-minute fixation using ice-cold 2% paraformaldehyde (PFA) in PB (pH 7.4). Following fixation, left and right liver lobe pieces, tail tip, perigonadal fat, and uterine tissue samples were collected, snap-frozen in liquid nitrogen, and stored at -80°C. Femur and tibia were collected from both the right and left hind limbs; muscle tissue was removed, and bones were stored in 4% PFA at 4 °C for 2 days, after which they were washed 6 times for 15 minutes in 0.01 M PBS, then finally stored in 70% ethanol at 4°C until micro-CT scan. Brains were collected after perfusion and stored in 2% PFA at 4°C overnight. The brains were transferred to 30% sucrose solution in 0.1 M PB the following day at 4°C, then once sunk (24-48 h), flash-frozen in hexane and stored at -80°C until further processing. Mouse bodies were stored at -20°C until body composition analysis was performed.

### Statistical analysis

All statistical analyses were performed in R (Version 4.4.2) and RStudio (Version 2024.12.1+563) [44,45]. Data cleaning, visualization, and analysis were performed using the tidyverse (v2.0.0) [46], ggplot2 (v3.5.1) [47], and ggpubr (v0.6.0) [48] packages. Linear mixed-effect models (LMMs) and generalized linear mixed-effect models (GLMMs) were fitted using the lme4 package (v1.1-35.5)[49] via lmer() and glmer() functions.

Fixed effects included parity (1-, 2-, 4 cycles of pregnancy and lactation or virgin), strain (BALB/cByJ or C57BL/6J), and their interactions. The inclusion of interaction terms were assessed using a likelihood-ratio test, comparing nested models with and without the interaction term on models fitted with maximum likelihood. Final models were refitted using restricted maximum likelihood. Batch (1–5) was included as a random effect. For repeated-measures outcomes, time (e.g., postpartum day, timepoint) was included as a fixed effect, and individual identity nested within batch was modeled as a random intercept (1 | Batch/ID). Body weight trajectories were additionally modeled using a third-order polynomial of postpartum day and litter size as a covariate to account for non-linear growth and nursing demand. In addition, food intake was modelled as a second-order polynomial of postpartum day and litter size as a covariate to account for non-linear rise in food intake during lactation. For non-normally distributed outcomes (i.e. latency outcomes in behavior tests), GLMMs or GLMs with Gamma distributions were applied as appropriate. For fecal corticosterone concentration, the outcome variable was log-transformed before fitting the linear model.

Model selection followed a hierarchical simplification approach in cases of singular fit or convergence issues, including reduction or removal of random effects and, when necessary, use of (general) linear models. Model assumptions (normality and homoscedasticity of residuals) were verified using residual visual diagnostics. Full model specifications, sample sizes, and fit statistics (R², ICC, AIC, log-likelihood, and F-statistics) for all outcome variables are provided in Table S3. Statistical significance of fixed effects was assessed using Satterthwaite’s approximation for degrees of freedom, as implemented in the lmerTest package (v3.1.3)[50], and model results were summarized and reported using the modelsummary (v2.1.1) [51] and kableExtra (v1.4.0) [52] packages. One-time pregnant BALB/cByJ dams were defined as the reference category (intercept), with fixed-effect coefficients interpreted as deviations from this baseline. Results are reported as estimated marginal means with 95% confidence intervals (CIs). All raw data, including R scripts and model outputs, are available on OSF (https://osf.io/8pqc6 [53]).

## Data, Materials, and Software Availability

All data and R scripts for the project are publicly available at: OSF (https://osf.io/8pqc6)

## Acknowledgements.

This work was supported by the National Research Program (NRP) Number 79 Advancing 3R – Animals, research and society sponsored by the Swiss National Science Foundation (SNSF) project number 206453 (to CNB), and a UFAW Animal Welfare Student Scholarship Award (to GS). Micro-CT imaging was performed with equipment maintained by the Swiss Center for Musculoskeletal Imaging, SCMI, Balgrist Campus AG, Zürich. We acknowledge and thank the many colleagues who supported this work through their technical assistance and discussions: Hanno Würbel, Thorsten Buch, Thomas A. Lutz, Madeleine Gries, Roman Stadler, Jonas Zaugg, Mohammed Hankir, Urs Meyer, Christine Nicol, and Josep Call.

