## Supplemental Material for "Evidence of maternal resilience in two mouse strains in the context of permanent mouse breeding strategies"

### Supplemental Results

***LDB.*** The reduction in distance travelled was not observed in the C57BL/6J strain ($\beta_{4 \times C57BL/6J}$ = 11.7, 95% CI [2.4, 21.0], p = 0.015). Similar tendencies were visible in the relative time spent in the light box (Fig. S2B).

***OFT.*** Similar trends were observed in time spent in the center zone with C57BL/6J spending longer in the center zone compared to the BALB/cByJ strain ($\beta_{C57BL/6J}$ = 218.6, 95% CI [152.7, 284.5], p < 0.001; $\beta_{BALB/cByJ}$ = 113.6, 95% CI [67.0, 160.2]) Fig. S2C). In addition, there was a significant interaction between parity levels and strains, suggesting a strain-specific response to pregnancy and lactation. Virgin animals spent longer in the center in the BALBc/ByJ strain ($\beta_{Virgin \times BALB/cByJ}$ = 77.5, 95% CI [11.6, 143.4], p = 0.022; $\beta_{1 \times BALB/cByJ}$ = 113.6, 95% CI [67.0, 160.2]), whereas there was a decrease in the C57BL/6J virgins ($\beta_{Virgin \times C57BL76J}$ = -124.8, 95% CI [ -218.1, -31.6], p = 0.009). Further, BALB/cByJ two-time pregnant dams did not show a significant change in time spent in the center, whereas C57BL/6J had a significant decrease ($\beta_{2 \times BALB/cByJ}$ = 49.5, 95% CI [-18.3, 117.2], p = 0.149; $\beta_{2 \times C57BL/6J}$ = -178.2, 95% CI [-274.0, -82.5], p < 0.001). A reduction in the total distance moved in four-time pregnant dams was not present in the OFT ($\beta_4$ = -1074.5cm, 95% CI [-2762.3, 613.4], p = 0.208; $\beta_1$ = 10457.9 cm, 95% CI [9287.0, 11628.7]), potentially because of the prolonged time to weaning (Fig. 3I).

***SI***. Nine to fifteen days after weaning of the last litter, dams were tested in the social interaction test. The number of pregnancy and lactation cycles did not affect interactions with either the prisoner or dummy mouse. However, a strong strain effect was observed, with C57BL/6J dams spending more time interacting with both the prisoner and the dummy mouse and moving overall less than BALB/cByJ dams (Fig. S2D-S2F).

***SPT***. Anhedonia-like behavior, assessed by a sucrose preference test around 30 days post-weaning, differed by strain but was unaffected by the reproductive experience. C57BL/6J animals had, on average, a 16.8% higher sucrose preference compared to the BALB/cByJ strain ($\beta_{C57BL/6J}$ = 16.8%, 95% CI [8.6, 24.9], p < 0.001; $\beta_{BALB/cByJ}$ = 58.3, 95% CI [48.9, 67.7]) (Fig. S2G).

***Oxytocin***. Oxytocin-positive cell counts in the PVN varied with reproductive history. To assess this, oxytocin immunoreactivity was quantified by immunohistochemistry five weeks after weaning of the final litter. Both four-time dams and virgin animals exhibited fewer oxytocin-positive cells compared to one-time dams across both strains ($\beta_4$ = -12.9, 95% CI [-23.5, -2.2], p = 0.019; $\beta_{Virgin}$ = -9.4, 95% CI [-18.6, -0.1], p = 0.048; $\beta_1$ = 27.5, 95% CI[17.2, 37.7]) (Fig. S2H).

### Supplemental Materials & Methods

**Pup retrieval assessment**

To assess maternal motivation, a five-minute pup retrieval test was conducted in the early dark phase (between 11 am and 1 pm) on PD3, PD6, and PD9. Each session was recorded using infrared webcams (Raspberry Pi NoIR camera RPi-CAM-V2 Night Vision connected to a Raspberry Pi 3 Model B+ board). Whenever possible, four pups (two females and two males) were removed from the litter and kept under a red warming light for 10 minutes. If fewer than four pups were in the litter, then all pups were removed. One minute before starting the test, the food grid in the home cage was removed, and the nest position was noted. Pups were returned to the two opposite corners of the nest, separated by male and female pups. The placement of the pups in the right-left corners was alternated by sex on each test day. Five minutes of each video were coded by an experimenter blinded to the dam's reproductive cycle using the Behavioral Observation Research Interactive Software (BORIS), a free, versatile open-source event-logging software for video coding and live observations [53]. Video analysis started once the experimenter's hands were completely off the cage. If the dam had already approached the pups when experimenters' hands were still in the cage, the moment the pups were added to the cage was considered the starting point. The following behavioral parameters were analyzed: latency to retrieve the first pup to the nest, latency to retrieve all pups to the nest, latency to sniff the first pup, total time spent sniffing the first pup, and time spent on the nest, divided into pre- and post-retrieval. An individual's behavior was initiated as soon as the behavior was observed. In the case of time spent on the nest, the beginning and the end of the behavior were set once the upper body of the dam was respectively inside and outside the nest. If the dam was seen building the nest, this was coded as time spent on the nest, even if the upper body of the dam was outside of the nest. The observer coding the videos was blinded to the dam's experimental group. To assess intra-observer reliability, the first three videos from each test day (P3, P6, and P9) were reevaluated after all videos were rated, achieving 95% agreement. Trials in which not all pups were returned to the nest were excluded from the latency analyses.

**Elevated plus maze test (EPM)**

An elevated plus maze test (EPM) was performed on PD24 ± 3 days (with one female tested 7 days after weaning) during the dark phase of the light cycle (11 am – 6 pm) in a room adjacent to the housing room. One two-time pregnant female C57BL/6J mouse was excluded from analysis because testing occurred five days outside the predefined postpartum testing window. The EPM was conducted in infrared light (< 1 lux) using a plus-shaped maze with 35.5 cm-long arms of which two had open walls, and two had arms enclosed with 15 cm-high opaque acrylic walls. Animals were introduced to the middle portion of the maze facing an open arm and were allowed to explore freely for 5 minutes. Time spent and distance traveled in the open and closed arms, center zone, and number of head dips were measured by an automated video-tracking system (Noldus Information Technology, The Netherlands). An entry into the central, open, or closed arm zone is detected when the central point of the subject's body crosses the border in the respective area, while a head dip is recorded when the subject's nose protrudes over the edge of the open arms into the head dip zone. The maze was cleaned with water and tissue between different subjects.

**Light – dark box test (LDB)**

A light-dark box test (LDB) was performed on PD27 ± 3 days during the dark phase of the light cycle (4 – 9 pm) in a nearby testing room. Mice were transported inside a lightproof metallic box with four compartments to the test chambers (30 x 30 x 24 cm; Multi Conditioning System, TSE Systems GmbH, Bad-Homburg, Germany). Each test chamber was divided into two compartments, one brightly lit area (100 lux) and one dark area (1 lux), separated by a dark Plexiglas wall and interconnected by a 3.5 x 10 cm gate. Mice were placed in the dark compartment and, following a 5-second habituation phase, gates between compartments were automatically opened, and the animals' location and movement across both chambers were tracked by a beam-break system for 10 minutes. The maze was cleaned with water and tissue between different animals.

**Open field test (OFT)**

An open field test (OFT) was performed on PD30 ± 3 days during the dark phase of the light cycle (11 am – 5 pm) in a directly adjacent room to the housing room illuminated by infrared light (< 1 lux) . One two-time pregnant female C57BL/6J mouse was excluded from analysis because testing occurred six days outside the predefined postpartum testing window. The open field consisted of four open arenas (51 x 51 cm) with 40 cm wall height, allowing simultaneous testing of up to four animals. Dams were always placed in the same corner of the maze, facing the open arena, and were allowed to explore freely for 30 minutes. Time spent in the center zone, time spent in the periphery, and total distance moved were measured by an automated video-tracking system (Noldus Information Technology, The Netherlands), in which a center zone (27.5 x 27.5 cm) and a peripheral zone around the central arena were designated. A transition between zones was detected when the central point of the subject's body crossed the border into the respective area. After each mouse, the maze was cleaned with water and tissue.

**Social interaction test (SI)**

A social interaction test was performed on PD33 ± 3 days during the dark phase of the light cycle (11 am – 5 pm) in a room directly adjacent to the housing room, illuminated by infrared light (< 1 lux). The Y-maze consisted of three equally sized arms (37.5 x 9 cm) extending from a central platform. Each arm was enclosed by a 10 cm high transparent Plexiglas wall and covered with a lid of the same material, leaving access to the central platform. Two of the arms are equipped with grid cages (13 x 8 x 10 cm) at the end of the arm, one containing an unfamiliar conspecific female mouse of the same strain (prisoner), the other one containing a small plush toy mouse matching the size and fur color of the strain (dummy). Animals were placed in the third, empty arm of the Y-maze and were allowed to explore for 5 minutes. Total distance moved was measured by an automated video-tracking system (Noldus Information Technology, The Netherlands). Due to some reflection on the plastic lid of the maze, interaction time with the prisoner or dummy mouse had to be evaluated by a blinded observer using BORIS [53]. An interaction was counted when the test animal is within the 2-cm zone directly in front of the wire grid, facing the prisoner or dummy with its snout.

**Oral glucose tolerance test (OGTT) and Insulin sensitivity test (IST)**

An oral glucose tolerance test was conducted on PD36 (+/- 3 days), followed by an insulin sensitivity test on PD40 (+/- 3 days), to assess the effect of repeated pregnancy and lactation on glucose homeostasis. Mice were fasted for 4h, starting 2h before dark onset. Baseline body weight and blood glucose were measured using Contour Next Blood Glucose Meter (Bayer Consumer Care AG, Basel, Switzerland), 30 minutes before the 4h fast was over. At the end of the 4h fasting period, mice received either an oral gavage of 50% glucose solution (5ml/kg body weight; 2.5g glucose/kg; B. Braun) for the OGTT or an intraperitoneal injection of insulin (0.5U/kg body weight) for the IST. Blood glucose was measured from the tail tip at -30, 15, 30, 60, 90, and 120 min relative to administration. In cases of hypoglycemia during the IST (blood glucose < 1.5 mmol/l accompanied by clinical signs such as tremor, apathy, or spasms), mice were immediately gavaged with 50% glucose solution. Fourteen mice required glucose administration, four were retested after a 4d recovery period. All fourteen animals were included in subsequent analyses. Food was returned after completion of the test. Area under the curve (AUC) was calculated from baseline using the trapezoidal rule.

**Indirect calorimetry**

Indirect calorimetry was performed with a 16-metabolic cage system (PhenoMaster; TSE systems, Bad Homburg, Germany) in the postweaning period, starting from PD42 (+/- 2 days). During this time, the dam's body weight was measured daily. Dams were acclimatized to the system for three days in groups of 2-3 animals, followed by two days of single housing. After this 5-day acclimatization phase, $O_2$ , and $CO_2$ levels, and food and water intake were measured by the TSE system for three consecutive days. $O_2$ consumption ($VO_2$) and $CO_2$ production ($VCO_2$) were calculated using the manufacturer's software (TSE PhenoMaster, version 6.0.1). Data related to $VO_2$ and $VCO_2$ were used to calculate energy expenditure (EE) and respiratory exchange ratio (RER) using Weir's equation (Weir, 1949) : *EE (kcal) = (3.941 × $VO_2$ + 1.106 × $VCO_2$)* / 1000. Data visualization and analysis were performed according to the consensus guide of Banks and colleagues [22]. Due to our 4x2 factorial study design (parity x strain), data analysis using CalR [54] was not possible, but equivalent analysis was performed using R. Measurements were acquired every 20 minutes per animal and averaged to hourly measurements. Continuous graphs display hourly measurements with mean $\pm$ SD, no rolling averages were used. Whenever possible, three-day averages were calculated for energy intake (kcal/h), RER, and EE (kcal/h). 10 animals had to be fully excluded from data analysis due to malfunctioning food hoppers, leading to a mismatch of energy intake and energy expenditure. In 7 animals an average of two days was calculated due to malfunctioning food hoppers on one measurement day, leading to the following group sizes (BALB/cByJ, / C57BL/6J: parity 1 (8/10), parity 2 (6/9), parity 4 (4/6), and virgin (8/10)).

**Sucrose preference test (SPT)**

Sucrose preference test (SPT) was conducted from days PD51 to PD53 (+/- 5 days), while dams were single-housed. A two-bottle choice test was performed over 2-3 days, and 23 hours of fluid intake was measured daily by weighing the bottles between 10-11 am. On day 0 of SPT, both bottles contained water for acclimatization and to assess the initial side preference of animals. On testing days 1-3 one bottle contained 1% sucrose solution and was alternately placed on the preferred or unpreferred side; water remained in the second bottle. The position of the two bottles was switched daily to distinguish between a sucrose or a side preference. Sucrose preference was calculated by the following formula: Sucrose preference (%) = sucrose consumed (g) / total liquid consumed *100. Mice were considered to have a side preference if they consumed 75% of the liquid from the same side on all three sucrose testing days, regardless of bottle content. Subjects demonstrating a side preference were excluded from the statistical analysis, resulting in the following group sizes (BALB/cByJ, / C57BL/6J: parity 1 (5/10), parity 2 (8/7), parity 4 (4/5), and virgin (8/8)).

**Body composition measurement using magnetic resonance imaging (EchoMRI)**

Post-mortem total lean and fat mass were measured around PD55 (± 5 days) using EchoMRI (Zurich Integrative Rodent Physiology, University of Zurich) [55]. Calibrations and measurements were performed according to the manufacturer's instructions. Two measurements were performed and averaged per mouse.

**Bone mass (micro-CT scans)**

Micro-computed tomography (micro-CT) scans were performed on the excised, defleshed tibia of breeding dams to assess trabecular bone mineral density and body length estimation using a micro-CT device (SkyScan 1176 In-Vivo Micro-CT, Bruker, Kontich, Belgium) at the Balgrist University Hospital Zurich in collaboration with Dr. Sander Botter. For analysis of trabecular and cortical bone structure, high-resolution scans of the proximal tibias were performed with the following scan parameters: 8 µm voxel size, 90kV source voltage, 278 µA source current, 0.3° rotation angle, 0.5 mm aluminum filter, 1000 ms exposure time, and frame averaging of 3. Images were reconstructed using NRecon V1.7.4.2 (Bruker/Skyscan) with the following settings: Smoothing: 2 (Gaussian), ring artifact correction 10, defect pixel mask 10, and beam hardening correction 20%. Following image reconstruction, the DataViewer software (Bruker/Skyscan) was used to rotate the images and align the long axis of the tibia consistently across all scanned bones. Bone density was calculated by defining a volume of interest (VOI) that extended from the end of the proximal tibial growth plate to a point 200 pixels distal to this starting location, encompassing the full tibial width. Analysis of trabeculae was conducted using CTAn V1.20.8.0 (Bruker/Skyscan). Mineralized structures were identified through global thresholding, with a lower threshold of 75 and an upper threshold of 255. The following parameters were calculated for all experimental subjects: Trabecular bone volume (BV), tissue volume (TV), Percentage bone volume (BV/TV), trabecular thickness (Tb.Th), trabecular number (Tb.N), trabecular separation (Tb.Sp), connectivity density (CD) and cortical bone thickness (Ct.Th). Tibia length was measured from a second, lower resolution scan. Whenever possible, the left tibia was scanned, unless the bone was damaged due to the extraction procedure, where the right tibia was analyzed instead. Voxel size was set to 35 µm with 90 kV source voltage, 278 µA source current, 1° rotation angle, and 360° scan, 0.5 mm aluminum filter, and 80 ms exposure time. After automated stitching of scans, images of tibias were rotated in the upright orientation using the software NRecon V1.7.4.2 and DataViewer (Bruker/Skyscan). The subsequent procedure and settings for reconstruction and 3D rotation were the same as in the high-resolution scans. The tibia length was calculated by counting the number of cross sections between the beginning of the tibia plateau (start of the tibia bone) and the medial malleolus (end of the tibia bone). The number of cross sections was then multiplied by the voxel size (0.035 mm) to calculate the total tibia length in mm.

**Fecal corticosterone metabolites measurement**

Fecal samples from the females were collected as previously described at three different time points during the mouse's lifespan: baseline sampling prior to mating, on the day of final weaning, and on the day of perfusion. Fecal samples were stored at -80 degrees until further processed. Corticosterone extraction was performed following the protocol established by Prof. Rupert Palme [17]. In brief, fecal samples were defrosted and dried using a drying cabinet (SalvisLab, Rotkreuz, Switzerland) at 70 degrees until fully desiccated. The dried samples were then ground and exactly 0.050 g (0.049-0.051 g) of feces was weighed into a new Eppendorf tube. If not enough feces was available, 0.045g was weighted into a new tube or until reaching the next possible 5 mg step. Subsequently, 80% methanol was added to the weighed samples, starting with 1 ml for 0.05 g and adjusting by deducting 0.1 ml for every 5 mg step down. The tubes were briefly hand-shaken to mix the contents and then placed in a thermo shaker (Labgene, Châtel-Saint-Denis, Switzerland) for 15 minutes at a frequency of 700 rpm at room temperature, followed by centrifugation for 10 minutes at 2500 G. Twice 0.25 ml of the supernatant were pipetted into clean tubes. The liquid phase of one tube was evaporated and sent to Vienna for measurement of corticosterone metabolites by a well established and validated enzyme immunoassay [17]. The other tube was stored at -80°C as a backup.

**Immunohistochemistry of brain tissue**
Frozen forebrains of experimental animals were sectioned in five series of 30 mm thickness on a cryostat (Leica CM3050 S; Biosystems; DE). Sections were mounted directly onto glass slides (Superfrost® Plus; Thermo Scientific; DE), beginning at the medial and caudal part of the hypothalamus, including the Paraventricular Nucleus (PVN) and the Arcuate Nucleus of the Hypothalamus (ARC) at -0.71 mm from bregma until -1.79 mm, according to the mouse brain atlas [56]. Slides were stored at -20°C in cryoprotectant solution (20% glycerol; 30% ethylene glycol; 50% phosphate buffer (0.1 M PB)) until staining. Brain sections were simultaneously stained for oxytocin and pSTAT3 expression using immunofluorescence protocols optimized for visualizing pSTAT3 and oxytocin [57,58]. Primary antibodies used were rabbit-α-pSTAT3 (1:500; Phopshp-Stat3 (Tyr705) (D3A7) XP, #9145 from Cell Signaling Technology) and mouse-α-Oxytocin (1:1000; Cat #MAB5296; EMD Millipore Corp, United States). Secondary antibodies used were donkey-α-rabbit–AlexaFluor647 and donkey-α-mouse–AlexaFluor488 (both 1:100; Jackson ImmunoResearch, Laboratories). Slides were stored at 4°C in the dark until they were scanned at the Center for Microscopy and Image Analysis, University of Zurich, Switzerland. Fluorescence images were taken at 20x magnification, with the coarse and fine focus set to the DAPI channel. Images were acquired with the following excitation parameters: AF405 (DAPI channel, LEDS 385 with 5.0% intensity, exposure time of 20 ms with 30% shift), AF488 (Oxytocin channel, LEDS 475 with 70% intensity, exposure time of 40 ms with 30% shift), AF555 (pSTAT3 channel, LEDS 555 with 100% intensity, exposure time of 300 ms with 30% shift). Fluorescent positive cells were automatically counted with QuPath v0.5.1 [59] within pre-defined regions of interest (ROIs) in the arcuate nucleus (bregma -1.43 until -1.91) for pSTAT3-positive cells and in the paraventricular nucleus (bregma -0.71 until -1.07) for oxytocin-expressing neurons. The following labeling thresholds were defined: p-STAT3 (Minimum area: 20 μm2, maximum area: 400 μm2, intensity threshold: 600) and oxytocin (Minimum area: 30μm2, maximum area: 400, intensity threshold: 1000). Left and right hemispheres were counted individually and summed up to the total number of cells per slide. Whenever possible, three slides per animal were averaged within the defined bregma areas.

**Gene expression analysis (qPCR)**
Gene expression analysis using quantitative polymerase chain reaction (qPCR) was performed to examine the expression of genes in liver and adipose tissue involved in fatty acid, glucose, and iron metabolism. In brief, RNA was extracted from frozen mouse adipose and liver tissue using the RNeasy® Lipid Tissue Mini Kit (QIAGEN AG, Germany, Cat. No. 74804) and following the manufacturers instructions. RNA concentration and purity were measured using the NanoDrop 2000 Spectrophotometer (Thermo Scientific, Switzerland, 25004826-0). Isolated RNA was reverse transcribed to cDNA using the SensiFAST cDNA Synthesis Kit (Meridian Bioscience, Cat. Nr. BIO-65053, BIO-65054), diluted 1:10 and stored at -20°C until further processing. SYBR Green reverse transcription (RT) PCR (FastStart Universal SYBR Green Master (ROX), Sigma Aldrich, Germany, Ref. 04913850001) was performed using the RT-PCR System (Applied Biosystems 7500, Center of Clinical Studies at the Vetsuisse Faculty of the University of Zurich). Each sample was analyzed in duplicates and the two strains were separated on two different plates.

Relative gene expression was determined using the ΔΔCt method: Cycle thresholds (Ct) were normalized against housekeeping genes (HK), Ribosomal protein L19 (RPL19) for adipose tissue and 36B4 for liver tissue, ΔCt = Ctgene – CtHK. Then each sample was compared to the mean of the control group (one-time pregnancy group in the respective strain), ΔΔCt = ΔCtsample – ΔCtcontrol, and the relative expression was calculated as 2-ΔΔCt. Primers were designed by Microsynth AG, Switzerland and diluted 1:10 with RNase-free water. For the sequences of the primers see Table below:

| Gene | Forward Sequences (5'-3') | Reverse Sequence (5'-3') |
|---|---|---|
| Ribosomal protein L19 (RPL19) | AAGCCTGTGACTGTCCATTC [1] | CTTCTTGGATTCCCGGTATC [1] |
| Ribosomal Protein Lateral Stalk Subunit P0 (RPLP0 / 36B4) | GGCCCTGCACTCTCGCTTTC [2] | TGCCAGGACGCGCTTGT [2] |
| Stearoyl-CoA Desaturase 1 (SCD1) | GCAAGCTCTACACCTGCCTCTT [3] | CGTGCCTTGTAAGTTCTGTGGC [3] |
| Fatty acid synthase (FASN) | CACAGTGCTCAAAGGACATGC [3] | CACCAGGTGTAGTGCCTTCCTC [3] |
| Six transmembrane epithelial antigen of prostate 4 (Steap4) | GGGAATCACTTCCTTGCCATCAG [3] | TCCGCCATACACCAAAGTGTGG [3] |
| Peroxisome proliferator-activated receptor alpha (Pparα) | ACCACTACGGAGTTCACGCATG [3] | GAATCTTGCAGCTCCGATCACAC [3] |
| Leptin (Lep) | GTCCAGGATGACACCAAAACC [4] | GACAAACTCAGAATGGGGTGAA [4] |
| Resistin | AGCGGATGAAGAACCTTTC [4] | GGAGGAGACTGTCCAGCAAT [4] |
| Fibroblast growth factor 21 (FGF21) | ACCTGGAGATCAGGGAGGAT [4] | CACCCAGGATTTGAATGACC [4] |
| Fatty acid binding protein 4 (Fabp4) | GACGACAGGAAGGTGAAGAG [5] | ACATTCCACCACCAGCTTGT [5] |
| Transferrin receptor 1 (Trf1) | GGCGCTTCCTAGTACTCCCT [6] | ACTTGCCGAGCAAGGCTAAA [6] |
| Ferroportin 1 (FPN1) | TGCAGGAGTCATTGCTGCTAG [6] | TGGAGTTCTGCACACCATTGAT [6] |
| Hepcidin (HAMP) | CAGGGCAGACATTGCGATAC [6] | TGCAACAGATACCACACTGGG [6] |
| Iron regulatory protein 1 (Irp1) | AGCCTTTGGGAGTGAACGC [6] | GATGACATGCTGCCTTTCCAC [6] |
| Glutathione peroxidase (GPX4) | GTGCATCCCGCGATGATTGGCG [6] | GATTACTTCCTGGCTCCTGCCTC [6] |
| Glucose-6-Phosphatase (G6p) | GGGCATCAATCTCCTCTGGG | AGATGACGTTCAAACACCGGA |
| Apolipoprotein B (ApoB) | TTAAAGGACTTTGGGACTGGC | TAGGGAGCCTAGCAATCTGG |
| Cholesterol 7-alpha-hydroxylase (CYP7A1 ) | AACGATACACTCTCCACCTTTG [7] | CTGCTTTCATTGCTTCAGGG [7] |

***Primer sequences Table***. *References: [1][60], [2][60], [3][61], [4][62], [5][63], [6][64], [7][65]*

**Colorimetric ferrozine-based assay**

Total iron concentration was measured in liver lysates by colorimetric ferrozine-based assay [66]. Homogenization of liver tissue was performed in NaOH lysis buffer using 1.4 mm ceramic beads and the Betin Precellys 24 Tissue Homogenizer. To normalize the iron levels, protein concentration from the lysate was determined by Bicinchoninic Acid protein assay (EMD Millipore Corp., Germany, LOT: 4003433, LOT: 4003435). Then, the iron concentration was determined following the previously described steps [67]. Samples were run in duplicate, and iron content was interpolated using a $FeCl_3$ standard curve. Iron concentrations were calculated using the standard curve, and the results were normalized to each sample's protein content.

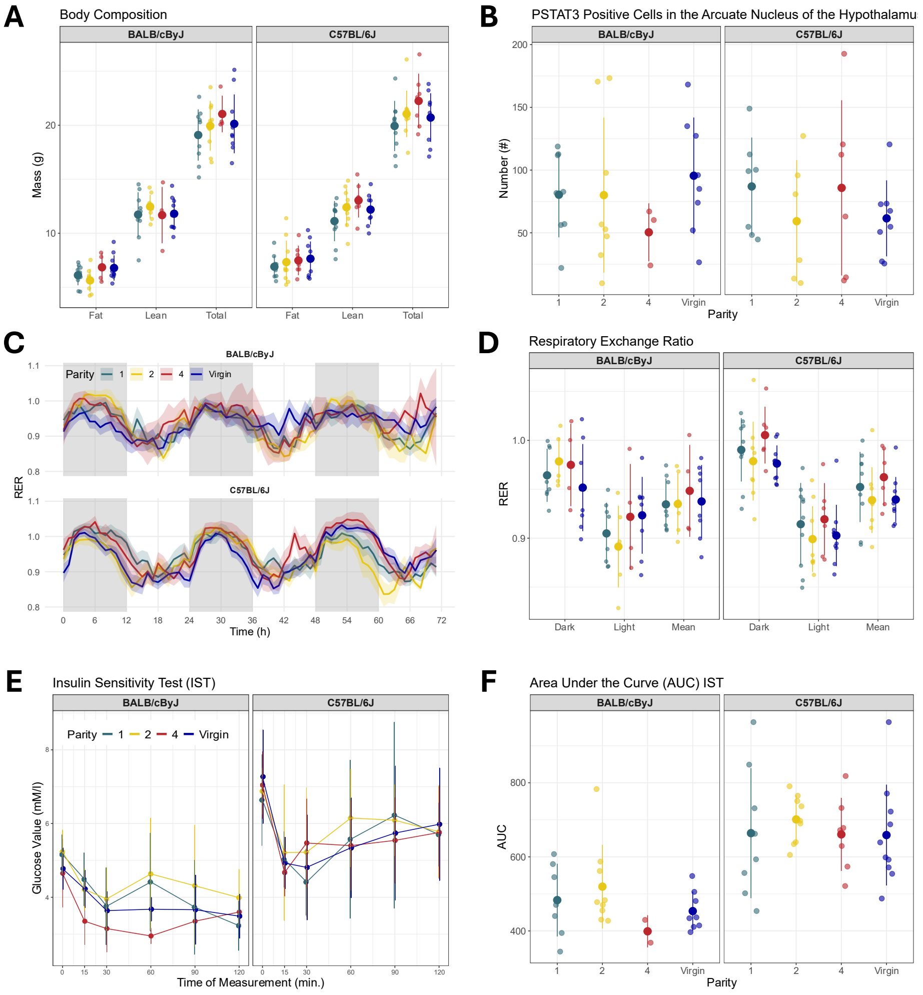

A
Body Composition
BALB/cByJ
C57BL/6J
Mass (g)
Fat
Lean
Total
B
PSTAT3 Positive Cells in the Arcuate Nucleus of the Hypothalamu
Number (#)
Parity
C
Parity 1 2 4 Virgin
RER
Time (h)
D
Respiratory Exchange Ratio
Dark
Light
Mean
E
Insulin Sensitivity Test (IST)
Glucose Value (mM/l)
Time of Measurement (min.)
F
Area Under the Curve (AUC) IST
AUC

Figure S1

**Figure S1**

Impact of reproductive load on body composition, leptin sensitivity, respiratory exchange ratio, and glucose homeostasis in the post-lactation period in female BALB/cByJ and C57BL/6J animals experiencing 1, 2, 4, or no (Virgin) cycles of concurrent pregnancy and lactation. (A) Fat, lean, and total mass of dams scanned with an Echo-MRI scanner to assess body composition after termination. (B) Number of phosphorylated signal transducer and activator of transcription factor 3 (PSTAT3) positive neurons in the arcuate nucleus of the hypothalamus after stimulation with 3 mg/kg leptin at termination. (C) Respiratory exchange ratio (RER) across three days from postpartum day (PD) 44-48. (D) Three-day RER during the dark phase, light phase, and their combined mean. (E) Blood glucose levels measured around PD 40 at 0, 15, 30, 60, 90, and 120 min following an intraperitoneal insulin (0.5U/kg body weight) injection administered after a 4-hour fast. (F) Area under the curve (AUC) for the insulin sensitivity test (IST). Data are represented as mean ± SD. Dots represent individual animals. Sample sizes for each outcome are provided in SI Appendix, Table S3.

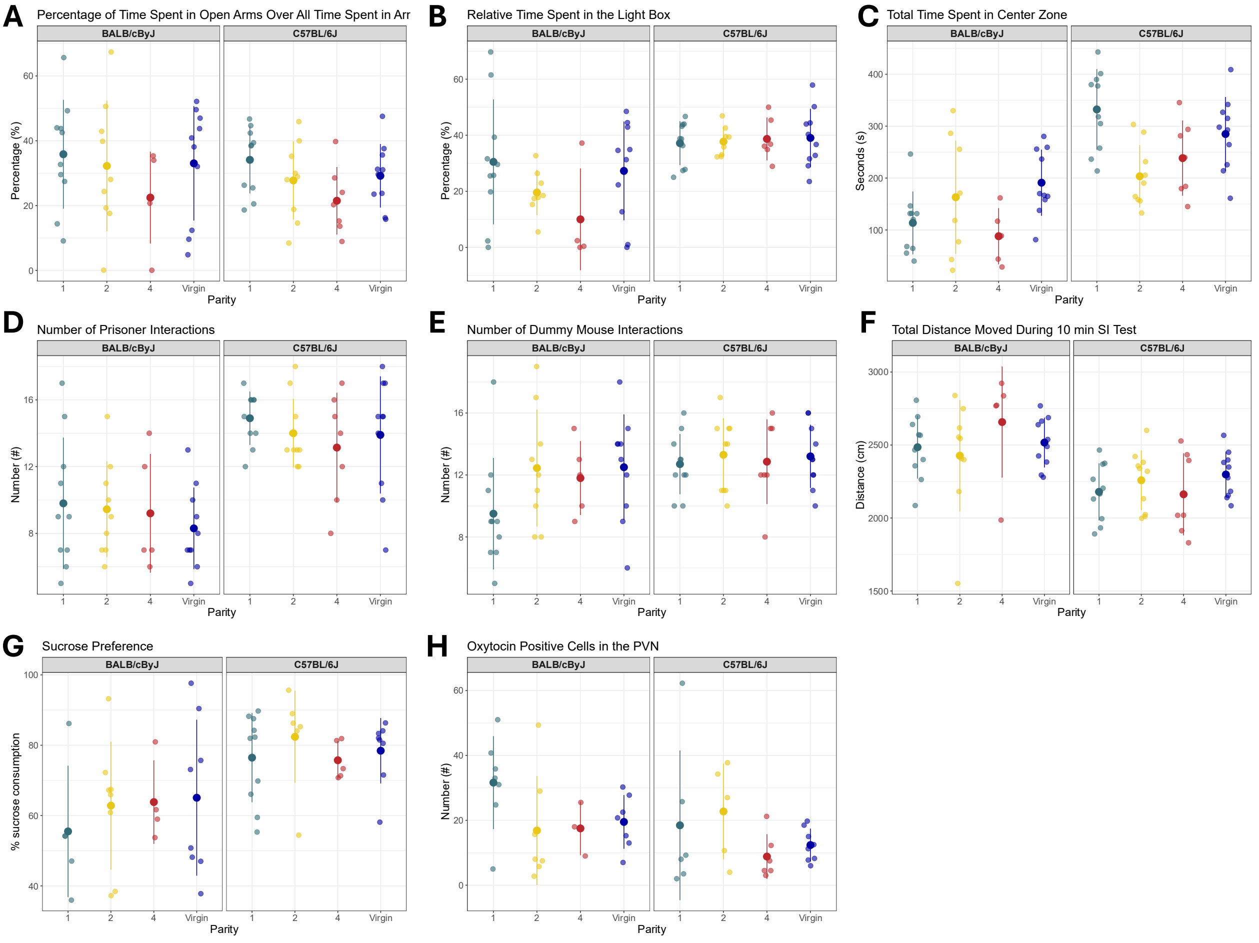

A
Percentage of Time Spent in Open Arms Over All Time Spent in Arr
B
Relative Time Spent in the Light Box
C
Total Time Spent in Center Zone
D
Number of Prisoner Interactions
E
Number of Dummy Mouse Interactions
F
Total Distance Moved During 10 min SI Test
G
Sucrose Preference
H
Oxytocin Positive Cells in the PVN
BALB/cByJ
C57BL/6J
Percentage (%)
Seconds (s)
Number (#)
Distance (cm)
% sucrose consumption
Parity
1
2
4
Virgin

Figure S2

**Figure S2**

Impact of reproductive load on maternal behavior, exploratory behavior, and locomotor activity. Female BALB/cByJ and C57BL/6J mice experienced 1, 2, or 4 cycles of pregnancy and lactation, or remained virgin (Virgin). (A) Percentage of time spent in the open arms during a 5-minute elevated plus maze test (EPM) conducted around postpartum day (PD) 24. (B) Relative time spent in the light box during a 10-minute light-dark-box (LDB) test paradigm run around PD27. (C) Total time spent in the center zone of a 30-minute open field (OF) test around PD30. (D) Number of prisoner and (E) dummy mouse interactions during a 10-minute social interaction (SI) test performed around PD33. (F) Total distance moved during the SI test. (G) Percentage of sucrose water consumption over all water consumed during a 36-hour sucrose preference test conducted from PD50-53. (H) Number of oxytocin-positive neurons in the paraventricular nucleus of the hypothalamus (PVN) after termination. Data are represented as mean ± SD. Dots represent individual animals. Sample sizes for each outcome are provided in SI Appendix, Table S3.

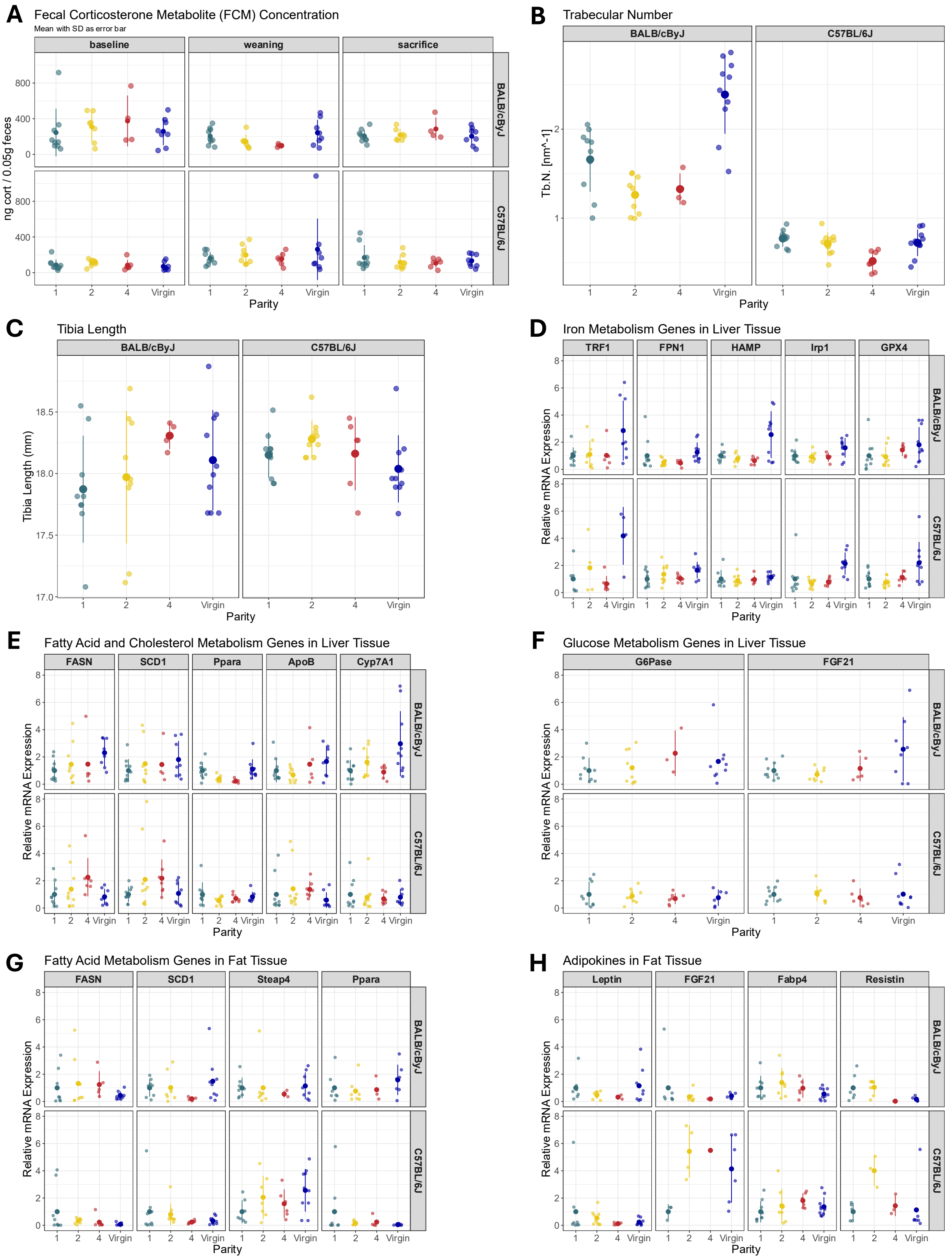
A
Fecal Corticosterone Metabolite (FCM) Concentration
Mean with SD as error bar
baseline
weaning
sacrifice
BALB/cByJ
C57BL/6J
ng cort / 0.05g feces
Parity
B
Trabecular Number
BALB/cByJ
C57BL/6J
Tb.N. [mm^-1]
Parity
C
Tibia Length
BALB/cByJ
C57BL/6J
Tibia Length (mm)
Parity
D
Iron Metabolism Genes in Liver Tissue
TRF1
FPN1
HAMP
Irp1
GPX4
Relative mRNA Expression
Parity
E
Fatty Acid and Cholesterol Metabolism Genes in Liver Tissue
FASN
SCD1
Ppara
ApoB
Cyp7A1
Relative mRNA Expression
Parity
F
Glucose Metabolism Genes in Liver Tissue
G6Pase
FGF21
Relative mRNA Expression
Parity
G
Fatty Acid Metabolism Genes in Fat Tissue
FASN
SCD1
Steap4
Ppara
Relative mRNA Expression
Parity
H
Adipokines in Fat Tissue
Leptin
FGF21
Fabp4
Resistin
Relative mRNA Expression
Parity

Figure S3

**Figure S3**

Impact of reproductive load on fecal corticosterone, bone structure, and genes involved in iron, fatty acid, cholesterol, and glucose metabolism in BALB/cByJ and C57BL/6J dams experiencing 1, 2, 4, or no (Virgin) cycles of concurrent pregnancy and lactation. (A) Absolute fecal corticosterone metabolites (FCMs) concentration in ng/0.05g feces before breeding onset (baseline), at weaning of the last litter (weaning), and on the termination day of the animals (sacrifice). (B) Trabecular number measured during microcomputer tomography (micro-CT) scan of the proximal tibia after killing. (C) Tibia length in millimeters measured on micro-CT scans of the full bone length. (D) Relative hepatic mRNA expression of genes involved in iron metabolism (TRF1, FPN1, HAMP, IRP1, GPX4). (E) Relative hepatic mRNA expression of genes involved in fatty acid and cholesterol metabolism (FASN, SCD1, Ppara, ApoB, Cyp7A1). (F) Relative hepatic mRNA expression of genes involved in glucose metabolism (G6Pase, FGF21). (G) Relative adipose tissue mRNA expression of genes involved in fatty acid metabolism (FASN, SCD1, Steap4, Ppara). (H) Relative adipose tissue mRNA expression of adipokine-encoding genes (Leptin, FGF21, Fabp4, Resistin). Data are represented as mean ± SD. Dots represent individual animals. Sample sizes for each outcome are provided in SI Appendix, Table S3. Abbreviations: TRF1, transferrin receptor 1; FPN1, ferroportin 1; HAMP, hepcidin; Irp1, iron regulatory protein 1; GPX4, glutathione peroxidase 4; FASN, fatty acid synthase; SCD1, stearoyl-CoA desaturase 1; Ppara, peroxisome proliferator-activated receptor alpha; ApoB, Apolipoprotein B; Cyp7A1, Cholesterol 7-alpha-hydroxylase; G6Pase, glucose-6-phosphatase; FGF21, fibroblast growth factor 21; Steap4, six-transmembrane epithelial antigen of prostate 4; Fabp4, fatty acid-binding protein 4.

**Table S1. Reproductive performance and offspring outcomes of BALB/cByJ and C57BL/6J dams across all consecutive breeding cycles.** Data are shown as mean ± SD (minimum, maximum) or percentage (n/N), where n indicates the number of animals meeting the specified criterion and N the total number of bred animals. Summary litter parameters across all litter per dam. PD, postnatal day.

| BALB/cByJ | One-time pregnant | Two-time pregnant | Four-time pregnant |
|---|---|---|---|
| Total number of pups born per dam | 7.1 ± 2 (5, 11) | 12 ± 3.16 (8, 17) | 18.8 ± 2.17 (17, 22) |
| Total number of pups weaned per dam | 5.5 ± 2.6 (0, 10) | 10.8 ± 3.5 (6, 16) | 18.2 ± 2.2 (17, 22) |
| Mean number of pups born across all litters per dam | 7.1 ± 2 (5, 11) | 6 ± 2.2 (2, 9) | 4.7 ± 2.7 (1, 9) |
| Mean number of pups weaned across all litters per dam | 5.5 ± 2.6 (0, 10) | 5.4 ± 2.7 (1, 9) | 4.6 ± 2.7 (1, 9) |
| Early pup mortality (PD1 / PD2) | 5.6% (N = 71/67) | 7.4% (N = 108/100) | 2.1% (N = 94/92) |
| Late pup mortality (PD2 / PD21) | 17.9% (N = 67/55) | 3% (N = 100/97) | 1.1% (N = 92/91) |
| Overall pup mortality (PD1 / PD21) | 22.5% (N = 71/55) | 10.2% (N = 108/97) | 3.2 % (N = 94/91) |

| C57BL/6J | One-time pregnant | Two-time pregnant | Four-time pregnant |
|---|---|---|---|
| Total number of pups born per dam | 8.2 ± 2 (4, 11) | 14.7 ± 2.8 (11, 19) | 28.4 ± 5 (22, 34) |
| Total number of pups weaned per dam | 7.8 ± 2 (4, 10) | 13.4 ± 2.1 (11, 16) | 25 ± 4.9 (17, 31) |
| Mean number of pups born across all litters per dam | 8.2 ± 2 (4, 11) | 7.4 ± 1.9 (3, 12) | 7.1 ± 2.6 (3, 13) |

| | | | |
|---|---|---|---|
| Mean number of pups weaned across all litters per dam | 7.8 ± 2 (4, 10) | 6.7 ± 1.8 (3, 10) | 6.3 ± 2.4 (1, 11) |
| Early pup mortality (PD1 / PD2) | 4.9% (N = 82/78) | 8.2% (N = 147/135) | 10.6% (N = 199/178) |
| Late pup mortality (PD2 / PD21) | 0% (N = 78/78) | 0.7% (N = 135/134) | 1.7% (N = 178/175) |
| Overall pup mortality (PD1 / PD21) | 4.9% (N = 82/78) | 8.8% (N =147/134) | 12.1% (N = 199/175) |

**Table S2**. Coefficients of the polynomial mixed-effects model. Cubic polynomial terms (intercept, x–$x^3$) are shown by strain (BALB/cByJ, C57BL/6J) and parity (1, 2, 4, Virgin). Negative cubic terms ($x^3$) indicate increasing trajectories followed by a decline, with larger absolute values reflecting greater curvature. Values closer to zero indicate a straighter curve. Predicted intercepts represent the sum of model components (Parity, strain, number of pups). Predicted max_y and max_x denote the peak body weight and the corresponding postpartum day, respectively.

| Strain Level | Pregnancy Level | Intercept | Strain | Pregnancy | Pups | x | x^2 | x^3 | Total_Intercept | Prediction Max_x | Prediction Max_y | Turning Point |
|---|---|---|---|---|---|---|---|---|---|---|---|---|
| BALB/cByJ | 1 | 29.28314 | 0 | 0 | 0.13918 | -0.43291 | 0.06888 | -0.00227 | 29.42232 | 16.297 | 31.132 | 1 |
| C57BL/6J | 1 | 29.28314 | -2.32986 | 0 | 0.13918 | 0.42128 | 0.00812 | -0.00087 | 27.09246 | 16.135 | 32.645 | 1 |
| BALB/cByJ | 2 | 29.28314 | 0 | 1.15619 | 0.13918 | -0.35738 | 0.04112 | -0.00126 | 30.57851 | 15.811 | 30.552 | 1 |
| C57BL/6J | 2 | 29.28314 | -2.61632 | 1.15619 | 0.13918 | 0.4772 | 0.00874 | -0.00106 | 27.96219 | 15.306 | 33.828 | 1 |
| BALB/cByJ | 4 | 29.28314 | 0 | 1.89489 | 0.13918 | 0.10135 | 0.00275 | -0.00019 | 31.3172 | 19.342 | 33.28 | 1 |
| C57BL/6J | 4 | 29.28314 | -1.76697 | 1.89489 | 0.13918 | 0.29745 | 0.02026 | -0.00128 | 29.55023 | 15.541 | 34.58 | 1 |
| BALB/cByJ | Virgin | 29.28314 | 0 | -2.93171 | 0 | 0.077 | -0.0004 | -0.00001 | 26.35143 | 21 | 28.151 | 0 |
| C57BL/6J | Virgin | 29.28314 | -1.33857 | -2.93171 | 0 | 0.23618 | -0.00905 | 0.00013 | 25.01286 | 21 | 27.674 | 0 |

**Table S3.** Statistical model specifications and fit indices for all presented analyses. Each row corresponds to a figure panel and reports the outcome variable, experimental design (group sizes), measurement, model type, fixed and random effects structure, number of observations, variance explained (marginal and conditional $R^2$ for linear mixed models [LMMs], or $R^2$ and adjusted $R^2$ for linear models), intraclass correlation coefficient (ICC), Akaike information criterion (AIC), and whether to use a model with or without interaction (likelihood ratio test [LRT] for mixed models or F-test for linear models). The R script filename generating each analysis is provided and matches script names available at https://osf.io/8pqc6. LMM, linear mixed model; LM = linear model; GLMM = generalized linear mixed model; ICC intraclass correlation coefficient; AIC, Akaike information criterion; LRT, likelihood ratio test; df, degrees of freedom.

<table>
<thead>
<tr><th>Figure</th><th>Outcome variable (unit)</th><th>Experimental design</th><th>Measurement</th><th>Model type</th><th>Fixed effects</th><th>Random effects</th><th>Number of Observations</th><th>$R^2$ (marg / cond or $R^2$ / $R^2$ adjusted)</th><th>ICC</th><th>AIC</th><th>LRT ($\chi^2$, df, p) or F-test (F, df1, df2, p)</th><th>Interaction retained</th><th>R file name</th></tr>
</thead>
<tbody>
<tr><td>Fig. 1C-D</td><td>body weight (g)</td><td>BALB/cByJ, C57BL/6J: parity 1 (10,10), parity 2 (9,10), parity 4 (5,7), and virgins (10,10)</td><td>every 3 days postpartum (3-21)</td><td>Gaussian LMM</td><td>Parity * strain * poly(time, 3) + pups</td><td>(1|batch/ID)</td><td>497</td><td>(0.53 / 0.95)</td><td>0.9</td><td>1532.8</td><td>$\chi^2$(25) = 159.5, p < 0.001</td><td>yes</td><td>2024_bw_analysis_v8_gramm_aim1.Rmd</td></tr>
<tr><td>Fig. 1E</td><td>body weight (g)</td><td>BALB/cByJ, C57BL/6J: parity 1 (10,10), parity 2 (9,10), parity 4 (5,7), and virgins (10,10)</td><td>on posptartum day 15 (highest weight gain)</td><td>Gaussian LMM</td><td>Parity + strain</td><td>(1|batch)</td><td>71</td><td>(0.55 / 0.58)</td><td>0.1</td><td>336.9</td><td>$\chi^2$(3) = 6.97, p = 0.07</td><td>no</td><td>2025_bw_PPD3_15_21_AL_v2_aim1.Rmd</td></tr>
<tr><td>Fig. 1F</td><td>body weight (g)</td><td>BALB/cByJ, C57BL/6J: parity 1 (10,10), parity 2 (9,10), parity 4 (4,7), and virgins (10,10)</td><td>at sacrifice (postpartum day 55 ± 5)</td><td>Gaussian LMM</td><td>Parity + strain</td><td>(1|batch)</td><td>70</td><td>(0.17 / 0.44)</td><td>0.3</td><td>293.1</td><td>$\chi^2$(3) = 1.27, p = 0.74</td><td>no</td><td>2024_body_weight_length_age_sacrifice_aim1.Rmd</td></tr>
<tr><td>Fig. 1G-H</td><td>food intake (kcal/h)</td><td>BALB/cByJ, C57BL/6J: parity 1 (10,10), parity 2 (9,10), parity 4 (5,7), and virgins (10,10)</td><td>every 3 days postpartum (3-21)</td><td>Gaussaian LMM</td><td>Parity * strain * poly(time, 2) + pups</td><td>(1|batch/ID)</td><td>464</td><td>(0.92 / 0.96)</td><td>0.4</td><td>-234.7</td><td>$\chi^2$(17) = 400.98, p < 0.001</td><td>yes</td><td>2025_last_lact_fi_AL_v2_cubic_aim1.Rmd</td></tr>
<tr><td></td><td></td><td></td><td></td><td></td><td></td><td></td><td></td><td></td><td></td><td></td><td></td><td></td><td></td></tr>
<tr><td>Fig. 2C</td><td>energy intake (kcal/h)</td><td>BALB/cByJ, C57BL/6J: parity 1 (8,10), parity 2 (6,9), parity 4 (4,6), and virgins (8,10) (animals excluded due to technical error, see supplementary methods for explanation)</td><td>average over 3d</td><td>Gaussian LMM</td><td>Parity + strain + mean bw 3d</td><td>(1|batch)</td><td>61</td><td>(0.16 / 0.21)</td><td>< 0.1</td><td>-92</td><td>$\chi^2$(3) = 2.36, p = 0.5</td><td>no</td><td>2025_TSE_CalR_copy_with_strain_v8_aim1_final_figures_manuscript.Rmd</td></tr>
<tr><td>Fig. 2C</td><td>energy intake (kcal/h)</td><td>BALB/cByJ, C57BL/6J: parity 1 (8,10), parity 2 (6,9), parity 4 (4,6), and virgins (8,10) (animals excluded due to technical error, see supplementary methods for explanation)</td><td>average over 3d during dark phase</td><td>Gaussian LMM</td><td>Parity + strain + mean bw 3d</td><td>(1|batch)</td><td>61</td><td>(0.23 / 0.37)</td><td>0.2</td><td>-26.1</td><td>$\chi^2$(3) = 4.35, p = 0.23</td><td>no</td><td>2025_TSE_CalR_copy_with_strain_v8_aim1_final_figures_manuscript.Rmd</td></tr>
<tr><td>Fig. 2C</td><td>energy intake (kcal/h)</td><td>BALB/cByJ, C57BL/6J: parity 1 (8,10), parity 2 (6,9), parity 4 (4,6), and virgins (8,10) (animals excluded due to technical error, see supplementary methods for explanation)</td><td>average over 3d during light phase</td><td>Gaussian LMM</td><td>Parity + strain + mean bw 3d</td><td>(1|batch)</td><td>61</td><td>(0.08 / 0.26)</td><td>0.2</td><td>-81.4</td><td>$\chi^2$(3) = 0.78, p = 0.85</td><td>no</td><td>2025_TSE_CalR_copy_with_strain_v8_aim1_final_figures_manuscript.Rmd</td></tr>
<tr><td>Fig. 2F</td><td>energy expenditure (kcal/h)</td><td>BALB/cByJ, C57BL/6J: parity 1 (8,10), parity 2 (6,9), parity 4 (4,6), and virgins (8,10) (animals excluded due to technical error, see supplementary methods for explanation)</td><td>average over 3d</td><td>Gaussian LMM</td><td>Parity + strain + mean bw 3d</td><td>(1|batch)</td><td>61</td><td>(0.39 / 0.58)</td><td>0.3</td><td>-196.8</td><td>$\chi^2$(3) = 0.95, p = 0.81</td><td>no</td><td>2025_TSE_CalR_copy_with_strain_v8_aim1_final_figures_manuscript.Rmd</td></tr>
<tr><td>Fig. 2F</td><td>energy expenditure (kcal/h)</td><td>BALB/cByJ, C57BL/6J: parity 1 (8,10), parity 2 (6,9), parity 4 (4,6), and virgins (8,10) (animals excluded due to technical error, see supplementary methods for explanation)</td><td>average over 3d during dark phase</td><td>Gaussian LMM</td><td>Parity + strain + mean bw 3d</td><td>(1|batch)</td><td>61</td><td>(0.45 / 0.61)</td><td>0.3</td><td>-164.2</td><td>$\chi^2$(3) = 1.34, p = 0.72</td><td>no</td><td>2025_TSE_CalR_copy_with_strain_v8_aim1_final_figures_manuscript.Rmd</td></tr>
<tr><td>Fig. 2F</td><td>energy expenditure (kcal/h)</td><td>BALB/cByJ, C57BL/6J: parity 1 (8,10), parity 2 (6,9), parity 4 (4,6), and virgins (8,10) (animals excluded due to technical error, see supplementary methods for explanation)</td><td>average over 3d during light phase</td><td>Gaussian LMM</td><td>Parity + strain + mean bw 3d</td><td>(1|batch)</td><td>61</td><td>(0.23 / 0.41)</td><td>0.2</td><td>-214.5</td><td>$\chi^2$(3) = 3.9, p = 0.27</td><td>no</td><td>2025_TSE_CalR_copy_with_strain_v8_aim1_final_figures_manuscript.Rmd</td></tr>
<tr><td>Fig. 2G</td><td>glucose (mM/l)</td><td>BALB/cByJ, C57BL/6J: parity 1 (10,10), parity 2 (9,10), parity 4 (5,7), and virgins (10,10)</td><td>blood glucose over time</td><td>Gaussian LMM</td><td>Parity * strain * timepoint</td><td>(1|batch/ID)</td><td>426</td><td>(0.82 / 0.82)</td><td>< 0.1</td><td>1660</td><td>$\chi^2$(38) = 90.02, p < 0.001</td><td>yes</td><td>2024_OGTT_v8_AL_aim1.Rmd</td></tr>
<tr><td>Fig. 2H</td><td>area under the curve (AUC)</td><td>BALB/cByJ, C57BL/6J: parity 1 (10,10), parity 2 (9,10), parity 4 (5,7), and virgins (10,10)</td><td>area under the curve (composite trapezoid rule)</td><td>Gaussian LMM</td><td>Parity * strain</td><td>(1|batch)</td><td>71</td><td>(0.80 / 0.81)</td><td>< 0.1</td><td>764.7</td><td>$\chi^2$(3) = 15.92, p < 0.001</td><td>yes</td><td>2024_OGTT_v8_AL_aim1.Rmd</td></tr>
<tr><td></td><td></td><td></td><td></td><td></td><td></td><td></td><td></td><td></td><td></td><td></td><td></td><td></td><td></td></tr>
<tr><td>Fig. 3A</td><td>seconds (s)</td><td>BALB/cByJ, C57BL/6J: parity 1 (10,10), parity 2 (9,10), and parity 4 (5,7)</td><td>latency to retrieve the first pup</td><td>Gamma GLMM (log link)</td><td>Parity + strain + PD</td><td>(1|ID)</td><td>144</td><td>(0.21 / 0.4)</td><td>0.2</td><td>1109.5</td><td>$\chi^2$(2) = 1.19, p = 0.55</td><td>no</td><td>pup_retrieval_v6_aim1.Rmd</td></tr>
<tr><td>Fig. 3B</td><td>seconds (s)</td><td>BALB/cByJ, C57BL/6J: parity 1 (10,10), parity 2 (7,10), and parity 4 (4,7)</td><td>latency to retrieve all pups</td><td>Gamma GLMM (log link)</td><td>Pairty + strain + PD + pups</td><td>(1|ID)</td><td>131</td><td>(0.27 / 0.52)</td><td>0.3</td><td>1292.5</td><td>$\chi^2$(2) = 1.72, p = 0.42</td><td>no</td><td>pup_retrieval_v6_aim1.Rmd</td></tr>
<tr><td>Fig. 3C</td><td>seconds (s)</td><td>BALB/cByJ, C57BL/6J: parity 1 (10,10), parity 2 (9,10), and parity 4 (5,7)</td><td>time spent on nest</td><td>Gaussian LM</td><td>Parity + strain + PD</td><td>na</td><td>147</td><td>(0.33 / 0.31)</td><td>na</td><td>1584.9</td><td>$F$(2, 139) = 2.91, p = 0.06</td><td>no</td><td>pup_retrieval_v6_aim1.Rmd</td></tr>
<tr><td>Fig. 3D</td><td>percentage (%)</td><td>BALB/cByJ, C57BL/6J: parity 1 (10,10), parity 2 (9,9), parity 4 (5,7), and virgins (10,10)</td><td>percentage of open arm entries</td><td>Gaussian LM</td><td>Parity + strain</td><td>na</td><td>70</td><td>(0.19 / 0.14)</td><td>na</td><td>525</td><td>$F$(3, 62) = 0.35, p = 0.79</td><td>no</td><td>EPM_v8_aim1.Rmd</td></tr>
<tr><td>Fig. 3E</td><td>number (#)</td><td>BALB/cByJ, C57BL/6J: parity 1 (10,10), parity 2 (8,8), parity 4 (4,6), and virgins (10,10)</td><td>number of light compartement entries</td><td>Gaussian LMM</td><td>Parity * strain</td><td>(1|batch)</td><td>66</td><td>(0.25 / 0.25)</td><td>< 0.1</td><td>407.3</td><td>$\chi^2$(3) = 8.53, p = 0.04</td><td>yes</td><td>LD_Box_v2_aim1.Rmd</td></tr>
<tr><td>Fig. 3F</td><td>number (#)</td><td>BALB/cByJ, C57BL/6J: parity 1 (10,10), parity 2 (9,9), parity 4 (5,7), and virgins (10,10)</td><td>number of center zone entries</td><td>Gaussian LM</td><td>Parity + strain</td><td>na</td><td>70</td><td>(0.32 / 0.28)</td><td>na</td><td>708.1</td><td>$F$(3, 62) = 1.01, p = 0.4</td><td>no</td><td>OFT_v3_aim1.Rmd</td></tr>
<tr><td>Fig. 3G</td><td>distance (cm)</td><td>BALB/cByJ, C57BL/6J: parity 1 (10,10), parity 2 (9,9), parity 4 (5,7), and virgins (10,10)</td><td>total distance moved during 5 min</td><td>Gaussian LM</td><td>Parity + strain</td><td>na</td><td>70</td><td>(0.56 / 0.54)</td><td>na</td><td>998</td><td>$F$(3, 62) = 0.01, p = 1</td><td>no</td><td>EPM_v8_aim1.Rmd</td></tr>
<tr><td>Fig. 3H</td><td>distance (cm)</td><td>BALB/cByJ, C57BL/6J: parity 1 (10,10), parity 2 (8,8), parity 4 (4,6), and virgins (10,10)</td><td>total distance moved during 10 min</td><td>Gaussian LMM</td><td>Parity + strain</td><td>(1|batch)</td><td>66</td><td>(0.62 / 0.6)</td><td>na</td><td>1154.4</td><td>$F$(3, 58) = 0.42, p = 0.74</td><td>no</td><td>LD_Box_v2_aim1.Rmd</td></tr>
<tr><td>Fig. 3I</td><td>distance (cm)</td><td>BALB/cByJ, C57BL/6J: parity 1 (10,10), parity 2 (9,9), parity 4 (5,7), and virgins (10,10)</td><td>total distance moved during 30 min</td><td>Gaussian LM</td><td>Parity + strain</td><td>na</td><td>70</td><td>(0.35 / 0.31)</td><td>na</td><td>1289.8</td><td>$F$(3, 62) = 1.15, p = 0.34</td><td>no</td><td>OFT_v3_aim1.Rmd</td></tr>
<tr><td></td><td></td><td></td><td></td><td></td><td></td><td></td><td></td><td></td><td></td><td></td><td></td><td></td><td></td></tr>
<tr><td>Fig. 4A</td><td>corticosterone (log-transformed)</td><td>BALB/cByJ, C57BL/6J: parity 1 (9,9), parity 2 (7,8), parity 4 (4,6), and virgins (8,8)</td><td>fecal corticosterone content (log-transformed)</td><td>Gaussian LMM</td><td>Parity + strain * timepoint</td><td>(1|ID)</td><td>177</td><td>(0.26 / 0.27)</td><td>< 0.1</td><td>372</td><td>$\chi^2$(2) = 19.45, p < 0.001</td><td>yes</td><td>corticosterone_analysis_v6_aim1.Rmd</td></tr>
<tr><td>Fig. 4C</td><td>percentage (%)</td><td>BALB/cByJ, C57BL/6J: parity 1 (10,10), parity 2 (9,10), parity 4 (4,7), and virgins (10,10)</td><td>bone volume / tissue volume (BV/TV in %)</td><td>Gaussian LM</td><td>Parity * strain</td><td>na</td><td>70</td><td>(0.86 / 0.85)</td><td>na</td><td>301.7</td><td>$F$(3, 62) = 166.49, p < 0.001</td><td>yes</td><td>BMD_inkl_20_slides_AL_v3_aim1.Rmd</td></tr>
<tr><td>Fig. 4D</td><td>micrometer (µm)</td><td>BALB/cByJ, C57BL/6J: parity 1 (10,10), parity 2 (9,10), parity 4 (4,7), and virgins (10,10)</td><td>trabecular thickness</td><td>Gaussian LM</td><td>Parity + strain</td><td>na</td><td>70</td><td>(0.36 / 0.32)</td><td>na</td><td>410.6</td><td>$F$(3, 62) = 1.04, p = 0.38</td><td>no</td><td>BMD_inkl_20_slides_AL_v3_aim1.Rmd</td></tr>
<tr><td>Fig. 4E</td><td>micrometer (µm)</td><td>BALB/cByJ, C57BL/6J: parity 1 (10,10), parity 2 (9,10), parity 4 (4,7), and virgins (10,10)</td><td>trabecular separation</td><td>Gaussian LM</td><td>Parity + strain</td><td>na</td><td>70</td><td>(0.78 / 0.76)</td><td>na</td><td>779.9</td><td>$F$(3, 62) = 1.86, p = 0.15</td><td>no</td><td>BMD_inkl_20_slides_AL_v3_aim1.Rmd</td></tr>
<tr><td>Fig. 4F</td><td>cubic millimeters ($mm^{-3}$)</td><td>BALB/cByJ, C57BL/6J: parity 1 (10,10), parity 2 (9,10), parity 4 (4,7), and virgins (10,10)</td><td>connectivity density</td><td>Gaussian LM</td><td>Parity * strain</td><td>na</td><td>70</td><td>(0.81 / 0.79)</td><td>na</td><td>567.4</td><td>$F$(3, 62) = 5.32, p < 0.001</td><td>yes</td><td>BMD_inkl_20_slides_AL_v3_aim1.Rmd</td></tr>
<tr><td>Fig. 4G</td><td>micrometer (µm)</td><td>BALB/cByJ, C57BL/6J: parity 1 (10,10), parity 2 (9,10), parity 4 (4,7), and virgins (10,10)</td><td>cortical bone thickness</td><td>Gaussian LM</td><td>Parity * strain</td><td>na</td><td>70</td><td>(0.87 / 0.85)</td><td>na</td><td>488.5</td><td>$F$(3, 62) = 3.54, p = 0.02</td><td>yes</td><td>BMD_inkl_20_slides_AL_v3_aim1.Rmd</td></tr>
<tr><td>Fig. 4H</td><td>micromol / mg protein</td><td>BALB/cByJ, C57BL/6J: parity 1 (10,10), parity 2 (9,10), parity 4 (5,7), and virgins (10,10)</td><td>iron concentration (umol/mg protein)</td><td>Gaussian LM</td><td>Parity + strain</td><td>na</td><td>71</td><td>(0.25 / 0.21)</td><td>na</td><td>294.9</td><td>$F$(3, 63) = 0.98, p = 0.41</td><td>no</td><td>Iron_concentration_R_v2_AL_aim1.Rmd</td></tr>
</tbody>
</table>

| | | | | | | | | | | | | | |
|---|---|---|---|---|---|---|---|---|---|---|---|---|---|
| Fig. S1A | mass (g) | BALB/cByJ, C57BL/6J: parity 1 (10,10), parity 2 (9,10), parity 4 (4,7), and virgins (10,10) | fat mass (g) | Gaussian LMM | Parity + strain | (1 \| batch) | 70 | (0.18 / 0.25) | 0.1 | 249.2 | $\chi^2$(3) = 2.03, p = 0.57 | no | 2026_body_composition_analysis_v9_aim1.Rmd |
| Fig. S1A | mass (g) | BALB/cByJ, C57BL/6J: parity 1 (10,10), parity 2 (9,10), parity 4 (4,7), and virgins (10,10) | lean mass (g) | Gaussian LMM | Parity + strain | (1 \| batch) | 70 | (0.07 / 0.47) | 0.4 | 261 | $\chi^2$(3) = 1.59, p = 0.66 | no | 2026_body_composition_analysis_v9_aim1.Rmd |
| Fig. S1A | mass (g) | BALB/cByJ, C57BL/6J: parity 1 (10,10), parity 2 (9,10), parity 4 (4,7), and virgins (10,10) | total mass (g) | Gaussian LMM | Parity + strain | (1 \| batch) | 70 | (0.13 / 0.27) | 0.2 | 317.6 | $\chi^2$(3) = 0.61, p = 0.89 | no | 2026_body_composition_analysis_v9_aim1.Rmd |
| Fig. S1B | number (#) | BALB/cByJ, C57BL/6J: parity 1 (8,7), parity 2 (8,6), parity 4 (3,6), and virgins (8,8) | number of pSTAT3 positive cells | Gaussian LMM | Parity + strain | (1 \| batch) | 54 | (0.01 / 0.19) | 0.2 | 540.7 | $\chi^2$(3) = 5.28, p = 0.15 | no | brain_R_script_v3_AL_aim1.Rmd |
| Fig. S1D | respiratory exchange ratio (RER) | BALB/cByJ, C57BL/6J: parity 1 (8,10), parity 2 (6,9), parity 4 (4,6), and virgins (8,10) (animals excluded due to technical error, see supplementary methods for explanation) | average over 3d | Gaussian LMM | Parity + strain | (1 \| batch) | 61 | (0.07 / 0.38) | 0.3 | -250.2 | $\chi^2$(3) = 1.27, p = 0.74 | no | 2025_TSE_CalR_copy_with_strain_v8_aim1_final_figures_manuscript.Rmd |
| Fig. S1D | respiratory exchange ratio (RER) | BALB/cByJ, C57BL/6J: parity 1 (8,10), parity 2 (6,9), parity 4 (4,6), and virgins (8,10) (animals excluded due to technical error, see supplementary methods for explanation) | average over 3d during dark phase | Gaussian LMM | Parity + strain | (1 \| batch) | 61 | (0.17 / 0.43) | 0.3 | -248.1 | $\chi^2$(3) = 2.24, p = 0.52 | no | 2025_TSE_CalR_copy_with_strain_v8_aim1_final_figures_manuscript.Rmd |
| Fig. S1D | respiratory exchange ratio (RER) | BALB/cByJ, C57BL/6J: parity 1 (8,10), parity 2 (6,9), parity 4 (4,6), and virgins (8,10) (animals excluded due to technical error, see supplementary methods for explanation) | average over 3d during light phase | Gaussian LMM | Parity + strain | (1 \| batch) | 61 | (0.04 / 0.32) | 0.3 | -227.3 | $\chi^2$(3) = 2.56, p = 0.46 | no | 2025_TSE_CalR_copy_with_strain_v8_aim1_final_figures_manuscript.Rmd |
| Fig. S1E | glucose (mM/l) | BALB/cByJ, C57BL/6J: parity 1 (7,8), parity 2 (9,9), parity 4 (2,7), and virgins (8,10) | blood glucose over time | Gaussian LMM | Parity * strain * timepoint | (1 \| batch/ID) | 360 | (0.40 / 0.60) | 0.3 | 1115.5 | $\chi^2$(38) = 47.98, p = 0.13 | no | 2025_IST_v4_AL_aim1.Rmd |
| Fig. S1F | area under the curve (AUC) | BALB/cByJ, C57BL/6J: parity 1 (7,8), parity 2 (9,9), parity 4 (2,7), and virgins (8,10) | area under the curve (composite trapezoid rule) | Gaussian LMM | Parity + strain | (1 \| batch) | 60 | (0.44 / 0.55) | 0.2 | 694.8 | $\chi^2$(3) = 0.36, p = 0.95 | no | 2025_IST_v4_AL_aim1.Rmd |
| Fig. S2A | percentage (%) | BALB/cByJ, C57BL/6J: parity 1 (10,10), parity 2 (9,9), parity 4 (5,7), and virgins (10,10) | percentage of time in open arms | Gaussian LM | Parity + strain | na | 70 | (0.1 / 0.04) | na | 576.7 | *F* (3, 62) = 0.05, p = 0.98 | no | EPM_v8_aim1.Rmd |
| Fig. S2B | percentage (%) | BALB/cByJ, C57BL/6J: parity 1 (10,10), parity 2 (8,8), parity 4 (4,6), and virgins (10,10) | percentage of time in light box | Gaussian LMM | Parity + strain | (1 \| batch) | 66 | (0.24 / 0.25) | < 0.1 | 521 | $\chi^2$(3) = 5.2, p = 0.16 | no | LD_Box_v2_aim1.Rmd |
| Fig. S2C | time (s) | BALB/cByJ, C57BL/6J: parity 1 (10,10), parity 2 (9,9), parity 4 (5,7), and virgins (10,10) | total time spent in the center zone | Gaussian LM | Parity * strain | na | 70 | (0.54 / 0.49) | na | 810.2 | *F* (3, 62) = 5.08, p < 0.001 | yes | OFT_v3_aim1.Rmd |
| Fig. S2D | number (#) | BALB/cByJ, C57BL/6J: parity 1 (10,10), parity 2 (9,10), parity 4 (5,7), and virgins (10,10) | number of prisoner interactions | Gaussian LM | Parity + strain | na | 71 | (0.44 / 0.41) | na | 359.3 | *F* (3, 63) = 0.23, p = 0.88 | no | social_interaction_v3_aim1.Rmd |
| Fig. S2E | number (#) | BALB/cByJ, C57BL/6J: parity 1 (10,10), parity 2 (9,10), parity 4 (5,7), and virgins (10,10) | number of dummy mouse interactions | Gaussian LM | Parity + strain | na | 71 | (0.13 / 0.08) | na | 358.2 | *F* (3, 63) = 0.81, p = 0.49 | no | social_interaction_v3_aim1.Rmd |
| Fig. S2F | distance (cm) | BALB/cByJ, C57BL/6J: parity 1 (10,10), parity 2 (9,10), parity 4 (5,7), and virgins (10,10) | total distance moved during 10 min | Gaussian LM | Parity + strain | na | 71 | (0.26 / 0.21) | na | 991.2 | *F* (3, 63) = 1.18, p = 0.33 | no | social_interaction_v3_aim1.Rmd |
| Fig. S2G | percentage (%) | BALB/cByJ, C57BL/6J: parity 1 (5,10), parity 2 (8,7), parity 4 (4,5), and virgins (8,8) | percentage sucrose consumption | Gaussian LM | Parity + strain | na | 55 | (0.26 / 0.20) | na | 459.3 | *F* (3, 47) = 0.27, p = 0.85 | no | sucrose_preference_v2_aim1.Rmd |
| Fig. S2H | number (#) | BALB/cByJ, C57BL/6J: parity 1 (7,6), parity 2 (7,5), parity 4 (3,6), and virgins (7,8) | number of oxytocin positive cells | Gaussian LMM | Parity + strain | (1 \| batch) | 49 | (0.14 / 0.35) | 0.2 | 375 | $\chi^2$(3) = 5.71, p = 0.13 | no | brain_R_script_v3_AL_aim1.Rmd |
| Fig. S3B | per nanometer ($nm^{-1}$) | BALB/cByJ, C57BL/6J: parity 1 (10,10), parity 2 (9,10), parity 4 (5,7), and virgins (10,10) | trabecular number | Gaussian LM | Parity * strain | na | 70 | (0.87 / 0.86) | na | 14.7 | *F* (3, 62) = 17.63, p < 0.001 | yes | BMD_inkl_20_slides_AL_v3_aim1.Rmd |
| Fig. S3C | milimeter (mm) | BALB/cByJ, C57BL/6J: parity 1 (9,10), parity 2 (9,10), parity 4 (4,6), and virgins (10,10) | tibia length (mm) | Gaussian LM | Parity + strain | na | 68 | (0.07 / 0.01) | na | 56.1 | *F* (3, 60) = 1.83, p = 0.15 | no | tibia_length_aim1.Rmd |
| Fig. S3D | relative mRNA expression | BALB/cByJ, C57BL/6J: parity 1 (10,6), parity 2 (9,4), parity 4 (5,7), and virgins (9,7) | transferrin receptor 1 (TRF1) | Gaussian LM | Parity * strain | na | 57 | (0.56 / 0.5) | na | 231.6 | *F* (3, 49) = 3.99, p = 0.01 | yes | Liver_iron_qPCR_v2_AL_aim1.Rmd |
| Fig. S3D | relative mRNA expression | BALB/cByJ, C57BL/6J: parity 1 (10,9), parity 2 (9,10), parity 4 (5,7), and virgins (10,9) | ferroportin 1 (FPN1) | Gaussian LMM | Parity + strain | (1 \| batch) | 69 | (0.2 / 0.21) | < 0.1 | 160 | $\chi^2$(3) = 4.27, p = 0.23 | no | Liver_iron_qPCR_v2_AL_aim1.Rmd |
| Fig. S3D | relative mRNA expression | BALB/cByJ, C57BL/6J: parity 1 (10,8), parity 2 (9,10), parity 4 (5,7), and virgins (10,9) | hepcidin (HAMP) | Gaussian LM | Parity * strain | na | 68 | (0.4 / 0.34) | na | 165.5 | *F* (3, 60) = 4.77, p < 0.001 | yes | Liver_iron_qPCR_v2_AL_aim1.Rmd |
| Fig. S3D | relative mRNA expression | BALB/cByJ, C57BL/6J: parity 1 (10,9), parity 2 (9,10), parity 4 (5,7), and virgins (10,9) | iron regulatory protein 1 (Irp1) | Gaussian LM | Parity + strain | na | 69 | (0.31 / 0.27) | na | 149.2 | *F* (3, 61) = 1.2, p = 0.32 | no | Liver_iron_qPCR_v2_AL_aim1.Rmd |
| Fig. S3D | relative mRNA expression | BALB/cByJ, C57BL/6J: parity 1 (10,9), parity 2 (9,10), parity 4 (5,7), and virgins (10,9) | glutathione peroxidase 4 (GPX4) | Gaussian LM | Parity + strain | (1 \| batch) | 69 | (0.22 / 0.25) | < 0.1 | 200.7 | $\chi^2$(3) = 1.54, p = 0.67 | no | Liver_iron_qPCR_v2_AL_aim1.Rmd |
| Fig. S3E | relative mRNA expression | BALB/cByJ, C57BL/6J: parity 1 (10,9), parity 2 (9,10), parity 4 (5,7), and virgins (9,9) | fatty acid synthase (FASN) | Gaussian LMM | Parity + strain | (1 \| batch) | 68 | (0.09 / 0.11) | < 0.1 | 319.8 | $\chi^2$(3) = 0.41, p = 0.94 | no | Liver_R_script_v2_AL_aim1.Rmd |
| Fig. S3E | relative mRNA expression | BALB/cByJ, C57BL/6J: parity 1 (10,9), parity 2 (9,10), parity 4 (5,7), and virgins (10,9) | stearoyl-CoA Desaturase-1 (SCD1) | Gaussian LMM | Parity + strain | (1 \| batch) | 69 | (0.08 / 0.09) | < 0.1 | 412.8 | $\chi^2$(3) = 0.67, p = 0.88 | no | Liver_R_script_v2_AL_aim1.Rmd |
| Fig. S3E | relative mRNA expression | BALB/cByJ, C57BL/6J: parity 1 (10,9), parity 2 (9,10), parity 4 (5,7), and virgins (10,9) | peroxisome proliferator-activated receptor alpha (Ppara) | Gaussian LMM | Parity + strain | (1 \| batch) | 69 | (0.2 / 0.21) | < 0.1 | 128.9 | $\chi^2$(3) = 4.37, p = 0.22 | no | Liver_R_script_v2_AL_aim1.Rmd |
| Fig. S3E | relative mRNA expression | BALB/cByJ, C57BL/6J: parity 1 (10,9), parity 2 (9,10), parity 4 (5,7), and virgins (8,9) | apolipoprotein B (ApoB) | Gaussian LMM | Parity + strain | (1 \| batch) | 67 | (0.02 / 0.2) | 0.2 | 213 | $\chi^2$(3) = 6.16, p = 0.1 | no | Liver_R_script_v2_AL_aim1.Rmd |
| Fig. S3E | relative mRNA expression | BALB/cByJ, C57BL/6J: parity 1 (10,9), parity 2 (9,10), parity 4 (5,7), and virgins (10,9) | cytochrom p450 7A1 (Cyp7A1) | Gaussian LM | Parity * strain | na | 69 | (0.3 / 0.22) | na | 232.9 | *F* (3, 61) = 2.9, p = 0.04 | yes | Liver_R_script_v2_AL_aim1.Rmd |
| Fig. S3F | relative mRNA expression | BALB/cByJ, C57BL/6J: parity 1 (9,9), parity 2 (9,10), parity 4 (5,7), and virgins (10,7) | glucose-6-phosphatase (G6Pase) | Gaussian LMM | Parity + strain | (1 \| batch) | 66 | (0.13 / 0.17) | 0.1 | 327.6 | $\chi^2$(3) = 5.49, p = 0.14 | no | Liver_R_script_v2_AL_aim1.Rmd |
| Fig. S3F | relative mRNA expression | BALB/cByJ, C57BL/6J: parity 1 (10,9), parity 2 (8,10), parity 4 (5,6), and virgins (9,9) | fibroblast growth factor 21 (FGF21) | Gaussian LM | Parity * strain | na | 66 | (0.29 / 0.21) | na | 241.6 | *F* (3, 58) = 2.9, p = 0.04 | yes | Liver_R_script_v2_AL_aim1.Rmd |
| Fig. S3G | relative mRNA expression | BALB/cByJ, C57BL/6J: parity 1 (8,10), parity 2 (8,9), parity 4 (5,7), and virgins (9,10) | fatty acid synthase (FASN) | Gaussian LMM | Parity + strain | (1 \| batch) | 65 | (0.08 / 0.16) | 0.1 | 246.9 | $\chi^2$(3) = 2.14, p = 0.54 | no | Markdown_fat_v4_AL_aim1.Rmd |
| Fig. S3G | relative mRNA expression | BALB/cByJ, C57BL/6J: parity 1 (8,10), parity 2 (8,9), parity 4 (5,7), and virgins (9,10) | stearoyl-CoA Desaturase-1 (SCD1) | Gaussian LMM | Parity + strain | (1 \| batch) | 65 | (0.01 / 0.13) | 0.1 | 243.2 | $\chi^2$(3) = 5.38, p = 0.15 | no | Markdown_fat_v4_AL_aim1.Rmd |
| Fig. S3G | relative mRNA expression | BALB/cByJ, C57BL/6J: parity 1 (8,9), parity 2 (8,9), parity 4 (3,6), and virgins (9,10) | six transmembrane epithelial antigen of prostate 4 (Steap4) | Gaussian LMM | Parity + strain | (1 \| batch) | 60 | (0.11 / 0.27) | 0.2 | 253.7 | $\chi^2$(3) = 5.61, p = 0.13 | no | Markdown_fat_v4_AL_aim1.Rmd |
| Fig. S3G | relative mRNA expression | BALB/cByJ, C57BL/6J: parity 1 (7,9), parity 2 (7,7), parity 4 (3,4), and virgins (8,9) | peroxisome proliferator-activated receptor alpha (Ppara) | Gaussian LMM | Parity + strain | (1 \| batch) | 53 | (0.11 / 0.17) | 0.1 | 250.6 | $\chi^2$(3) = 5.49, p = 0.14 | no | Markdown_fat_v4_AL_aim1.Rmd |
| Fig. S3H | relative mRNA expression | BALB/cByJ, C57BL/6J: parity 1 (7,8), parity 2 (8,9), parity 4 (3,6), and virgins (8,10) | leptin | Gaussian LMM | Parity + strain | (1 \| batch) | 59 | (0.08 / 0.1) | < 0.1 | 181.7 | $\chi^2$(3) = 2.89, p = 0.41 | no | Markdown_fat_v4_AL_aim1.Rmd |
| Fig. S3H | relative mRNA expression | BALB/cByJ, C57BL/6J: parity 1 (8,3), parity 2 (6,4), parity 4 (1,3), and virgins (5,7) | fibroblast growth factor 21 (FGF21) | Gaussian LM | Parity * strain | na | 33 | (0.7 / 0.62) | na | 134.2 | *F* (3, 25) = 5.27, p = 0.01 | yes | Markdown_fat_v4_AL_aim1.Rmd |
| Fig. S3H | relative mRNA expression | BALB/cByJ, C57BL/6J: parity 1 (8,9), parity 2 (8,9), parity 4 (5,7), and virgins (10,10) | fatty acid binding protein 4 (Fabp4) | Gaussian LM | Parity + strain | na | 66 | (0.13 / 0.07) | na | 310.6 | *F* (3, 58) = 0.98, p = 0.41 | no | Markdown_fat_v4_AL_aim1.Rmd |
| Fig. S3H | relative mRNA expression | BALB/cByJ, C57BL/6J: parity 1 (5,7), parity 2 (7,5), parity 4 (1,4), and virgins (5,8) | resistin | Gaussian LM | Parity + strain | na | 39 | (0.34 / 0.26) | na | 199 | *F* (3, 31) = 2.88, p = 0.05 | no | Markdown_fat_v4_AL_aim1.Rmd |
